# A novel reporter mouse for astrocyte-derived extracellular vesicles reveals trafficking of cargo to neuronal mitochondria

**DOI:** 10.64898/2026.04.16.718987

**Authors:** Xiaojia Ren, Zainuddin Quadri, Zhihui Zhu, Xu Fu, Liping Zhang, Erhard Bieberich

## Abstract

Extracellular vesicles (EVs) mediate intercellular transfer of lipids, proteins, and nucleic acids between nearly all cell types. We previously showed that astrocyte-derived EVs modulate neuronal mitochondria *in vitro*, but whether endogenous astrocytic EVs are trafficked to neuronal mitochondria *in vivo* remained unknown. To address this, we generated an EV reporter mouse, Aldh1l1-Cre; CD9-tGFP^fl/fl^, in which astrocyte-secreted EVs are labeled with a CD9-turboGFP fusion protein (CD9-tGFP). Astrocyte-specific expression of CD9-tGFP was verified in brain tissue and isolated EVs, comprising 13.2 ± 1.6% of total brain EVs. In primary glial cultures, CD9-tGFP was restricted to astrocytes, localizing to vesicular compartments and cell protrusions (filopodia and cilia), with 89.3 ± 2.2% of astrocyte-derived EVs carrying the label. These EVs were enriched with the sphingolipid ceramide, consistent with its co-distribution with CD9-tGFP in astrocytic cell protrusions. In the cortex, hippocampus, and cerebellum, CD9-tGFP was predominantly detected in astrocytic processes co-labeled with GLAST1 and GFAP, forming contacts with laminin-positive capillaries and parvalbumin-positive neurons. CD9-tGFP-labeled EVs were detected inside capillaries and neurons, and super-resolution STED microscopy revealed partial overlap with neuronal mitochondria. Live-cell spinning disk confocal imaging and AI-assisted proximity analysis confirmed uptake of CD9-tGFP EVs by neuronal cells and trafficking of their cargo to mitochondria *in vitro*. Biochemical isolation of synaptic and non-synaptic mitochondria confirmed EV-derived cargo on mitochondria *in vivo*, with 3-fold higher association of CD9-tGFP with synaptic than non-synaptic mitochondria. Together, these findings validate the Aldh1l1-Cre; CD9-tGFP^fl/fl^ reporter mouse as a powerful tool for tracking astrocyte-derived EVs *in vivo* and provide direct evidence that their cargo is preferentially trafficked to synaptic mitochondria.

**Graphical Abstract:** Astrocyte-derived extracellular vesicles target neuronal mitochondria *in vivo*

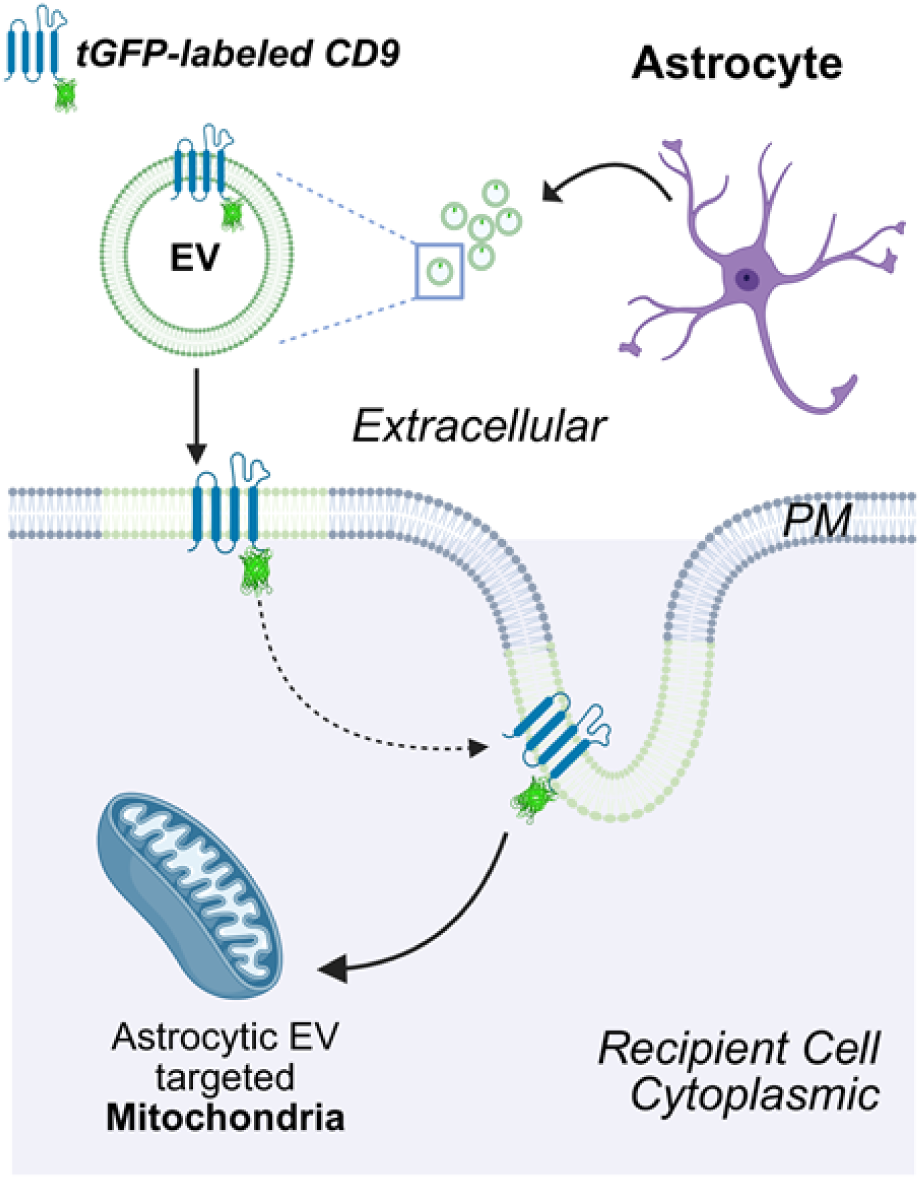

## Introduction

Extracellular vesicles (EVs) are a heterogeneous population of lipid bilayer-enclosed vesicles secreted by nearly all cell types. EVs mainly function as critical vectors for intercellular communication by transporting diverse cargo biomolecules such as lipids, proteins, nucleic acids and metabolites [1,2]. In the past two decades, a rapidly increasing number of studies focused on the essential roles of EVs in a plethora of biological processes, ranging from normal development and maintaining homeostasis to tissue repair, disease progression and pharmacological therapy [3–7]. In the central nervous system (CNS), EVs are critical for the intercellular communication among various cell types, including astrocytes, neurons, and microglia, influencing crucial processes such as synaptic plasticity, myelination, and the inflammatory response. EVs have been shown to mediate the transport and uptake of pathogenic factors in neurodegenerative diseases, resulting in the vulnerability of target cells and organelles [7–9]. On the other hand, studies have suggested the beneficial effects of EVs by enhancing the clearance of neurotoxic proteins, lipids and other pathogenetic factors [10,11].

In a blend of different types of EVs generated by a variety of tissues and cell types, often transported over large distances, it remains largely unclear which donor cells produce EVs and which target cells are affected by these EVs and how. These questions are fundamental to our understanding of the origin, fate, and function of EVs in neuronal disease. Most studies are conducted *in vitro* using cultured cells and analyzing EV composition without being able to track EVs *in vivo*. The lack of *in vivo* information is a severe obstacle to our understanding of the spatiotemporal dynamic and function of specific EVs populations, and to determine the physiological and pathological roles of EVs [12,13]. For example, we previously showed that astrocyte-derived EVs can modulate neuronal mitochondria *in vitro*, but whether endogenous astrocytic EVs target neuronal mitochondria *in vivo* remained unclear [14–16].

To address these critical gaps, we have developed a novel analytical platform for the generation and analysis of cell-type-specific EVs, which is based on a recently developed CD9-tGFP^fl/fl^ EV reporter mouse [17], which we crossbred with the Aldh1l1-Cre mouse [18], enabling tissue-specific excision of floxed genomic segments in astrocytes. In this model, astrocytes express CD9-turboGFP (CD9-tGFP), resulting in fluorescent labeling of astrocyte-derived EVs and enabling direct tracking of their origin, secretion, and fate *in vivo* and *in vitro*.

In this study, we characterize and validate this astrocyte-specific EV reporter mouse. Confocal imaging demonstrates that astrocytes release CD9-tGFP-labeled EVs that localize to capillaries and neurons. Using high-resolution live-cell spinning-disk confocal and super-resolution STED microscopy, combined with biochemical isolation of synaptic and non-synaptic mitochondria and AI-assisted proximity analysis, we show the association of CD9-tGFP-labeled cargo from astrocyte-derived EVs with neuronal mitochondria. In endothelial cells, EVs are taken up as well, however, EV cargo follows a bimodal intracellular distribution. Notably, we identify parvalbumin-positive cerebellar interneurons as major neuronal targets of astrocyte EVs, highlighting the ability of this reporter system to dissect cell-type- and organelle-specific EV interactions within the brain.

## Materials and Methods

### Animals

The CD9-tGFP^fl/fl^ mice [B6;129S1-*Gt(ROSA)26Sor^tm1(CAG-CD9/GFP)Dmfel^*/J] and Aldh1l1-Cre transgenic mice [B6;FVB-Tg(Aldh1l1-cre)JD1884Htz/J] were obtained from The Jackson Laboratory and subsequently bred in-house. The CD9-tGFP^fl/fl^ strain carries a loxP-flanked stop cassette inserted into the ROSA26 locus that allows Cre-dependent expression of a CD9-turboGFP fusion protein, enabling fluorescent labeling of EVs. The Aldh1l1-Cre line expresses Cre recombinase under control of the mouse Aldh1l1 promoter, allowing astrocyte-specific excision of floxed genomic segments. These two strains were crossed to generate Aldh1l1-Cre; CD9-tGFP^fl/fl^ mice following the breeding strategy illustrated in Fig. 1A.

**Figure 1.**
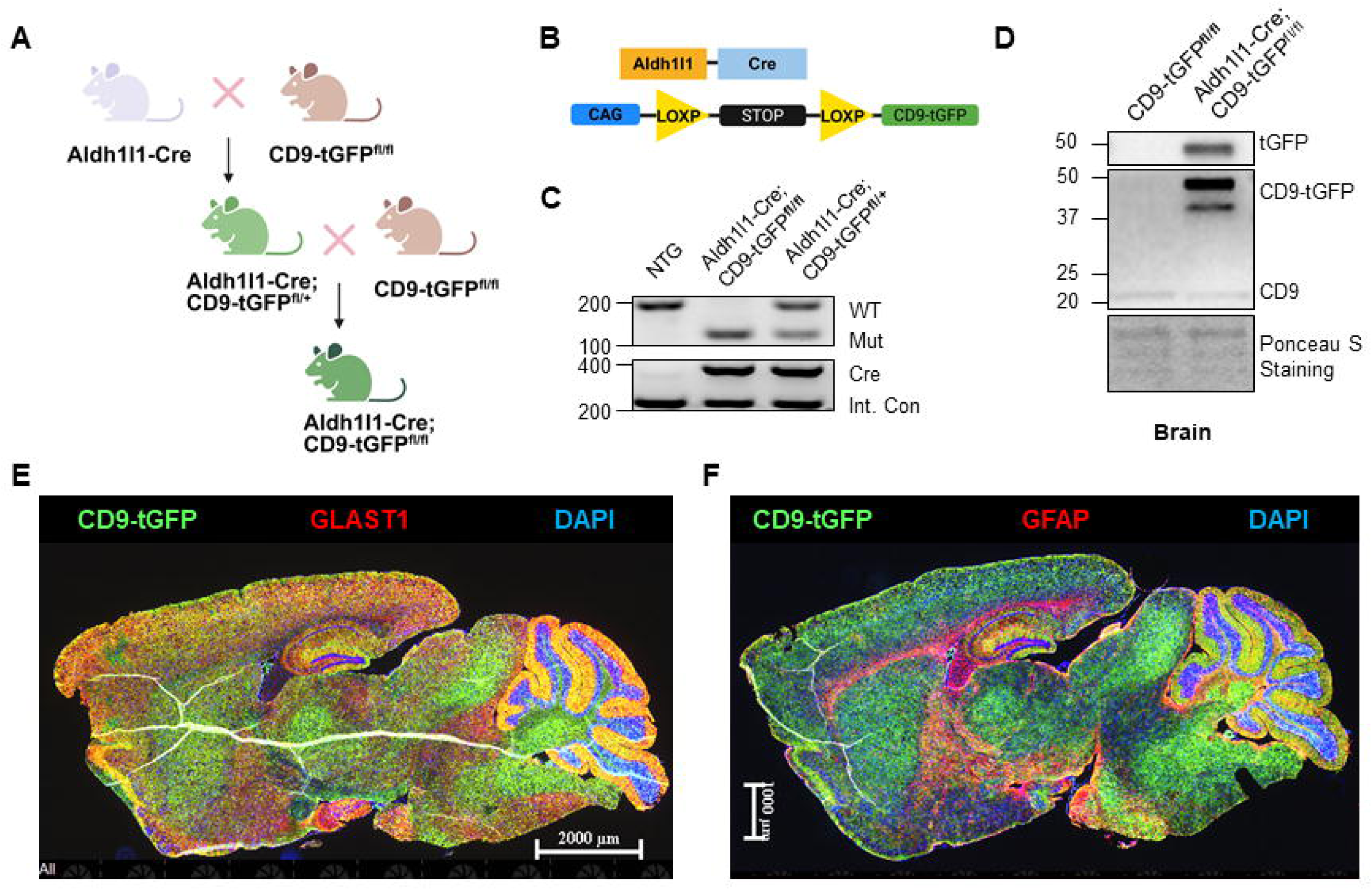
Generation of the reporter mouse and regional expression of astrocyte-specific CD9-tGFP fusion protein. (A) Breeding strategy for generating the Aldh1l1-Cre; CD9-tGFP^fl/fl^ mouse model. (B) Genetic schematic. Aldh1l1-Cre recombinase recognizes the LoxP sites and mediates the excision of the STOP cassette, leading to the astrocyte-specific expression of the CD9-tGFP fusion protein. (C) Genotyping results confirm successful excision of the STOP cassette in both heterozygous (Aldh1l1-Cre; CD9-tGFP^fl/+^) and homozygous (Aldh1l1-Cre; CD9-tGFP^fl/fl^) mice. (D) Immunoblots of brain lysates show expression of the CD9-tGFP fusion protein (around 50 kDa), detected by anti-CD9 and anti-tGFP antibodies, respectively; expression occurs exclusively in Cre-positive mice, alongside endogenous CD9 (around 25 kDa) present in all genotypes. (E, F) Immunofluorescence imaging of mouse brain sections co-stained for the astrocytic markers GLAST1 (E, red) and GFAP (F, red), respectively, with CD9-tGFP (green), showing that the green fluorescent signal colocalized with astrocytic markers throughout the mouse brain, particularly enriched in in hippocampus and cerebellum. Nuclei are stained with DAPI (blue).

Animals were group-housed in individually ventilated cages under a 12-h light/dark cycle with food and water provided *ad libitum*. Ear-punch biopsies were collected for genotyping. At the experimental endpoint, mice were anesthetized by isoflurane inhalation and perfused transcardially with ice-cold PBS. Brains were rapidly dissected on an ice-cooled metal stage and processed for protein extraction, EV isolation, mitochondrial isolation, or fixation for histological analysis. All animal procedures were performed in accordance with an approved Animal Use Protocol from the Institutional Animal Care and Use Committee (IACUC) at the University of Kentucky.

### Fixation and Cryosection of Mouse Brain

Perfused mouse brains were post-fixed in 4% paraformaldehyde (PFA) in PBS with gentle orbital shaking at 4 °C for at least one week. Following fixation, the PFA solution was discarded, and the brains were gently blotted dry with Kimwipes before immersion in 30% sucrose in PBS for cryoprotection. Brains were kept at 4 °C on an orbital shaker for 1-2 days until they sank to the bottom of the solution, indicating complete infiltration. Cryoprotected tissues were embedded in Tissue-Tek® O.C.T. compound (Sakura Finetek) over dry ice and stored at −80 °C. Sections were cut at the desired thickness using a Leica CM1860 cryostat and stored at −20 °C until immunohistochemical labeling.

### Isolation of EVs from Mouse Brain

Brain-derived EVs were isolated following a previously described protocol [14] with minor modifications (Suppl. Fig. 1A). After transcardial perfusion, whole brains were collected and cut into eight sagittal slices using a sterile scalpel in a Petri dish. The slices were transferred to Miltenyi Biotec C Tubes and processed using the Adult Brain Dissociation Kit (Miltenyi Biotec) according to the manufacturer’s instructions. Briefly, enzymatic dissociation buffer was added to the tissue in the C Tubes, and the “37C_ABDK_01” program on the gentleMACS™ Octo Dissociator (Miltenyi Biotec) was run to generate brain homogenates. The homogenate was passed through a 70-µm MACS SmartStrainer and rinsed with cold DPBS to obtain a final flow-through volume of approximately 10 mL.

The flow-through was centrifuged at 300 × *g* for 10 min at 4 °C, and the supernatant was supplemented with Halt™ Protease and Phosphatase Inhibitor Cocktail (Thermo Fisher Scientific, 1:100 dilution). Sequential centrifugation steps were then performed at 2,000 × *g* for 20 min and 10,000 × *g* for 30 min at 4 °C. The resulting supernatant was filtered through a 0.45-µm syringe filter and processed using the exoEasy Maxi Kit (Qiagen) following the manufacturer’s protocol. Briefly, samples were mixed with XBP binding buffer and loaded onto exoEasy spin columns, followed by centrifugation at 500 × *g* for 1 min at room temperature. Columns were washed with XWP buffer and centrifuged at 500 × *g* for 5 min. EVs were eluted with elution buffer and collected in the eluate for downstream analyses.

### Primary Mixed Glial Cell Culture

Mixed primary glial cells were isolated from postnatal day 3-4 (P3-P4) mouse pups following a previously published protocol [19]. Briefly, neonatal brains were dissected in cold Dulbecco’s modified Eagle’s medium (DMEM) supplemented with 100 U/mL penicillin-streptomycin (P/S), and meninges were carefully removed. The tissue was digested in DMEM containing 0.25% trypsin for 20 min at 37 °C, then gently triturated and passed through a 70-µm cell strainer to obtain a single-cell suspension. The suspension was centrifuged at 300 × *g* for 5 min, and the resulting cell pellet was resuspended in DMEM supplemented with 10% heat-inactivated fetal bovine serum (FBS) and 100 U/mL P/S. Cells were seeded into T-25 flasks and maintained at 37 °C in a humidified 5% CO_2_ incubator. Once cultures reached approximately 90% confluency, cells were detached using trypsin and either re-plated into T-75 flasks for expansion or onto poly-L-lysine-coated coverslips in 24-well plates for shaking and immunocytochemical experiments. Prior to experiments, cells were maintained overnight in phenol red-free DMEM without FBS.

### Isolation of Primary Astrocyte-Derived EVs

When primary astrocyte cultures (derived as described above) reached ∼90% confluency in T-75 flasks, the growth medium was replaced three times with phenol red-free, serum-free DMEM, each followed by a 10-min incubation at 37 °C to remove residual FBS. The flasks were then placed on an orbital shaker inside a humidified 37 °C, 5% CO_2_ incubator and shaken at 300 rpm for 24 h to promote EV release. After shaking, the conditioned medium was collected and centrifuged at 300 × *g* for 3 min at 4 °C to remove cellular debris. The supernatant was supplemented with Halt™ Protease and Phosphatase Inhibitor Cocktail (Thermo Fisher Scientific; 1:100 dilution) and sequentially centrifuged at 4,000 × *g* for 20 min, 10,000 × *g* for 30 min, and finally ultracentrifuged at 100,000 × *g* for 2 h at 4 °C. The resulting invisible pellet, containing astrocyte-derived EVs, was resuspended in DPBS. EVs were characterized by nanoparticle tracking analysis (NTA) using the ZetaView® system (Particle Metrix) and stored at −20 °C for subsequent experiments, including neuronal treatments and immunoblotting. The workflow is illustrated in Suppl. Fig. 1A.

### Neuro-2a (N2a) Cell Culture and Treatment with Astrocyte-Derived EVs

Mouse neuroblastoma N2a cells were cultured in Dulbecco’s modified Eagle’s medium (DMEM) supplemented with 10% heat-inactivated fetal bovine serum (FBS) and 100 U/mL penicillin-streptomycin (P/S) at 37 °C in a humidified incubator with 5% CO_2_. Cells were seeded either on coverslips in 24-well plates for immunocytochemical staining or in 24-well glass-bottom plates for live-cell imaging. Upon reaching confluency, cultures were rinsed with phenol red-free, serum-free DMEM and incubated overnight in the same medium for serum deprivation.

For immunolabeling, EVs were added to N2a cells at a concentration of 8000 EVs per cell and incubated for the indicated durations. Cells on coverslips were fixed in 4% paraformaldehyde (PFA) in PBS and stored at 4 °C until further processing.

For live-cell imaging, mitochondria were labeled with 500 nM MitoTracker™ Red CM-H₂XRos (Invitrogen) for 45 min at 37 °C, followed by washing with phenol red-free, serum-free medium to remove excess dye. Plasma membranes were labeled with 100 nM MemGlow™ 640 (Cytoskeleton Inc.) prepared in the same medium, incubated for 5 min at 37 °C, and then washed. The glass-bottom plates were mounted on a Nikon CSU-W1 SoRa spinning-disk confocal microscope. EVs were subsequently added to the N2a cells at a concentration of 10 EVs per cell, and time-lapse imaging was initiated immediately.

### Immunocytochemistry

Immunolabeling was performed as described previously [20–22]. In brief, cryosections were air-dried at room temperature for 15 min, circumscribed with a ImmEdge™ hydrophobic barrier pen (Vector Laboratories Inc.), and allowed to dry completely. Sections were rehydrated in PBS for 30 min, permeabilized with 0.2% Triton X-100 for 15 min, washed three times with PBS, and blocked in 3% ovalbumin in PBS for 1 h at 37 °C. Identical permeabilization, washing, and blocking steps were applied to cells cultured on coverslips. After blocking, samples were incubated overnight at 4 °C with primary antibodies diluted in 0.1% ovalbumin/PBS. The following day, samples were washed three times with PBS and incubated with fluorophore-conjugated secondary antibodies (prepared in 0.1% ovalbumin/PBS) for 2 h at 37 °C. After three final PBS washes, sections and coverslips were mounted with Fluoroshield™ containing DAPI (Sigma-Aldrich) to counterstain nuclei. Fluorescence imaging was performed using a Nikon Eclipse Ti2-E inverted microscope, a Nikon AXR inverted confocal microscope, or a Leica STELLARIS STED (stimulated emission depletion) super-resolution microscope. Images were processed and analyzed using Nikon NIS-Elements software equipped with a 3D deconvolution module and AI-assisted analysis after conversion to tiff images. Brain-derived and primary astrocyte-derived EVs were incubated with 5 µM Vybrant CM-DiI in PBS for 2 h at 37 °C. Free dye was removed by ultracentrifugation at 100,000 x g. The labeled EVs were resuspended in PBS, mixed 1:1 with mounting medium, and mounted with coverslips for fluorescence microscopy.

### Purkinje Cell EV Quantification

Cerebellar cryosections from Aldh1l1-Cre; CD9-tGFP^fl/fl^ mice and littermate controls (lacking Cre-mediated CD9-tGFP expression) were immunolabeled with anti-parvalbumin antibody to identify Purkinje cell soma and visualized for CD9-tGFP to label astrocyte-derived EVs, as described in the Methods section on immunocytochemistry. Images were acquired as described and calibrated using scale bars. Multi-panel images were split into individual channels for analysis. Purkinje cell soma were automatically detected from the parvalbumin channel using intensity thresholding (optimized per image) followed by morphological filtering to remove artifacts. Detected soma were manually verified and selected for analysis as indicated in figure legends. CD9-tGFP-positive particles within each soma were quantified by intensity thresholding with size filtering to exclude noise and debris. Particles were assigned to soma based on centroid localization within the soma mask. Results were expressed as particles per cell and particle density (particles/μm² soma area). Data are expressed as mean ± SEM. Correlations between particle number or particle density and soma area were assessed by Pearson correlation. All analyses were performed using custom Python scripts (NumPy, SciPy, scikit-image) with interactive parameter optimization in Google Colab; The analysis code will be available upon request publicly archived upon acceptance.

### EV-Mitochondria Apposition Analysis

N2a and primary mouse endothelial cells were treated with astrocyte-derived EVs as described in Methods (8000 particles/cell, 2-24 h). Images of CD9-tGFP signals and Tom20-labeled mitochondria were acquired from five random fields per condition after deconvolution and Z-projection. Custom Python code segmented EVs and mitochondria by intensity thresholding, then calculated edge-to-edge distances (minimum boundary pixel separation; 0 nm = contact) and centroid-to-centroid distances. Apposition was defined at contact (0 nm), ≤100, ≤200, ≤500, and ≤1000 nm. Monte Carlo randomization (1,000 iterations per image) validated non-random EV distribution by comparing observed to random particle placements within the cellular region of interest while preserving mitochondrial network geometry (p < 0.05). For binned distance enrichment analysis, 10,000 iterations were used. Groups were compared using Mann-Whitney U test. Bimodality of edge-to-edge distance distributions was assessed using the bimodality coefficient (BC > 0.555 indicates bimodal distribution) and Gaussian mixture models comparing 1-component vs. 2-component fits using Bayesian Information Criterion. Analysis used NumPy, SciPy, scikit-image, and scikit-learn. The analysis code will be publicly archived upon acceptance.

### Immunoblot Analysis

Equal amounts of protein from brain tissue lysates, isolated mitochondria, or EVs were mixed with 2× Laemmli sample buffer (Sigma-Aldrich), heated at 98 °C for 5 min, and cooled on ice. Samples were separated by SDS-PAGE using Bio-Rad TGX™ gradient precast gels and transferred to PVDF membranes (Millipore) at 100 V for 1 h. Membranes were blocked with 5% non-fat dry milk in Tris-buffered saline containing 0.1% Tween-20 (TBST) for 1 h at room temperature and incubated overnight at 4 °C with primary antibodies diluted in 1% non-fat dry milk/TBST. The next day, membranes were washed three times with TBST and incubated with HRP-conjugated secondary antibodies for 2 h at 37 °C. Immunoreactive bands were visualized using SuperSignal™ West Femto or West Pico chemiluminescent substrates (Thermo Fisher Scientific) and imaged with an Azure 300 chemiluminescent imaging system (Azure Biosystems).

### Isolation of Non-synaptic and Synaptic Mitochondria

Pure synaptic and non-synaptic mitochondria were isolated from mouse brain tissue following previously described protocols with minor changes [23,24] and as illustrated in Suppl. Fig. 1B. Briefly, brain tissues were homogenized in ice-cold mitochondrial isolation buffer containing 215 mM mannitol, 75 mM sucrose, 0.1% bovine serum albumin (BSA), 20 mM HEPES, and 1 mM EGTA (pH 7.2, adjusted with KOH). The homogenate was centrifuged at 1,300 × *g* for 3 min at 4 °C to remove cellular debris. The resulting supernatant was centrifuged at 13,000 × *g* for 10 min at 4 °C to obtain a crude mitochondrial pellet, which was resuspended in mitochondrial isolation buffer.

Crude mitochondria were loaded onto a discontinuous Ficoll gradient consisting of 7.5% (top) and 10% (bottom) layers and ultracentrifuged at 100,000 × *g* for 30 min at 4 °C. The pellet fraction was collected as non-synaptic mitochondria. The synaptosomal fraction at the interphase was carefully collected and disrupted using a nitrogen cavitation chamber at 1,200 psi for 10 min at 4 °C. The disrupted synaptosomes were then purified by centrifugation on a second 7.5%/10% Ficoll discontinuous gradient at 100,000 × *g* for 30 min at 4 °C, and the resulting bottom pellet was collected as the synaptic mitochondrial fraction.

## Results

### CD9-tGFP is Regionally Expressed in the Astrocyte-Specific Reporter Mouse

To generate the fluorescent EV reporter mouse, we crossbred an astrocyte-specific Cre driver line (Aldh1l1-Cre) with the floxed-stop CD9-turboGFP (CD9-tGFP) reporter mouse previously developed by the Feliciano laboratory [17]. The breeding scheme is shown in Fig. 1A. CD9 belongs to the tetraspanin family regulating multiple biological processes such as cell adhesion and migration, and has been recognized as biomarker for EVs and filopodia [25–27]. As depicted in Fig. 1B and verified by genotyping (Fig. 1C), Aldh1l1- Cre mediates removal of the stop codon leading to CD9-tGFP expression in astrocytes. Immunoblotting using brain lysate showed that without Cre expression, only endogenous CD9 (∼25 kDa) was detected. With Cre, an additional protein of 50 kDa was immunolabeled with antibodies against both CD9 and tGFP, verifying expression of the CD9-tGFP fusion protein exclusively in our crossbred astrocytic EV reporter mouse (Fig. 1D).

To verify model specificity, we performed immunocytochemistry on brain cryosections from mice with and without Cre. CD9-tGFP^fl/fl^ mouse brain sections lacking Cre showed only weak background fluorescence (Suppl. Fig. 2A). In Aldh1l1-Cre; CD9-tGFP^fl/fl^ mice, Cre recombination drove robust astrocytic CD9-tGFP expression. Co-labeling of CD9-tGFP with GLAST1 (Fig. 1E, individual channels in Suppl. Fig. 2B) and GFAP (Fig. 1F, individual channels in Suppl. Fig. 2C) varied by region; strong overlap with astrocytic markers was found in the neocortex, hippocampus, and cerebellum. In contrast, the brainstem and choroid plexus displayed bright CD9-tGFP labeling with little GLAST1 co-localization, consistent with Aldh1l1-driven recombination in perivascular glial populations associated with the dense capillary network in these regions.

### CD9-tGFP-positive EVs Carry Signatures of Membrane Protrusions and Microvesicles

To test for EV secretion, we isolated EVs from brain tissue and primary cultured astrocytes following protocols previously developed in our laboratory [14,27] (see schematics in Suppl. Fig. 1A). Brain EVs size and concentration were quantified by NTA (Fig. 2A) and the size distribution were consistent with previous analyses of EVs from wild-type mouse brain [14]. Immunoblotting of EVs from Aldh1l1-Cre; CD9-tGFP^fl/fl^ brains detected a ∼50 kDa CD9-tGFP fusion protein recognized by both anti-CD9 and anti-tGFP antibodies (Fig. 2B). As expected, brain-derived EVs contained established vesicular marker flotillin-2. To quantify the proportion of CD9-tGFP labeled EVs, we labeled total brain EVs with Vybrant-CM-diI and counted particles co-labeled for tGFP. Figure 2C (left orange bar) shows that 13.2% ± 1.6% of brain EVs were tGFP positive, indicating that this proportion of EVs was derived from astrocytes.

**Figure 2.**
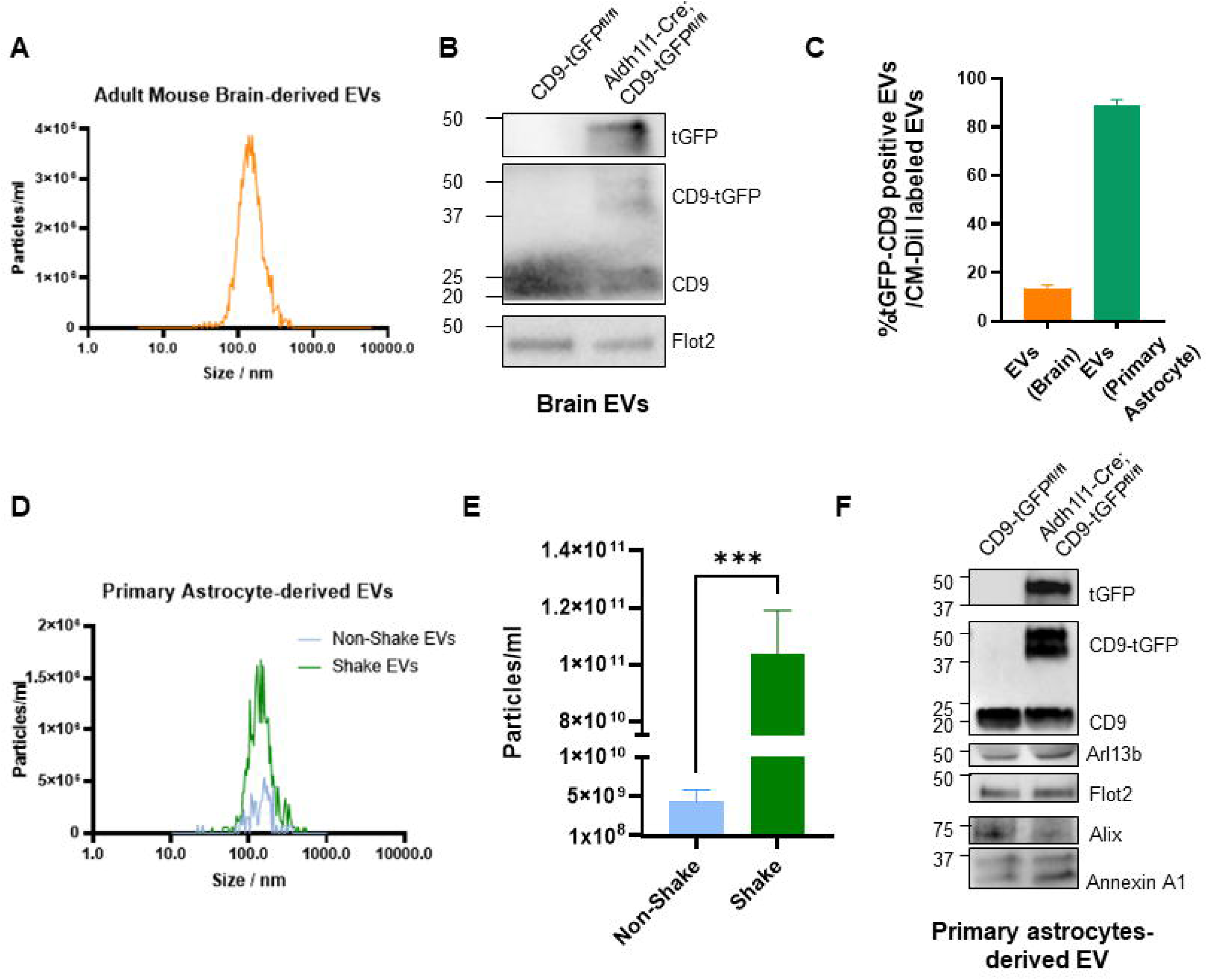
Characterization of EVs isolated from brain and primary astrocytes of the Aldh1l1-Cre; CD9-tGFP^fl/fl^ mouse model. (A) Representative NTA analysis of EVs isolated from brain tissue shows a particle size distribution and concentration within the expected EV range. (B) Immunoblots of brain-derived EVs confirm the presence of the CD9-tGFP fusion protein, detected by anti-CD9 and anti-tGFP antibodies, together with the EV marker Flotillin-2. (C) Quantification of the percentage of CD9-tGFP-positive EVs in brain-derived and primary astrocyte-derived EVs (stained with CM-Dil dye). (D) Representative NTA analysis of EVs isolated from primary astrocyte culture medium under static conditions (non-shaking) and after orbital shaking (shaking) shows a comparable size distribution between conditions. (E) Quantification of EV counts demonstrates that orbital shaking of primary cultured astrocytes increases EV yields compared to the non-shaking group (N=5, normalized to totally cell numbers, data are presented as mean ± SEM, t test ***p<0.001). (F) Representative immunoblots of primary astrocyte-derived EVs confirm the presence of CD9-tGFP, EV markers (Flotillin-2 and Alix), the microvesicle marker (Annexin A1), and the ciliary marker (Arl13b).

Mechanical stimulation has been shown to elevate EV secretion from other cell types such as mesenchymal stem cells, HeLa cells and dental pulp stem cells [28–31]. To increase our EV yield from primary astrocyte cultures, we applied an orbital shaking step originally introduced to remove microglia from mixed glial preparations [32–35], showing that mechanical stimulation enhances EV release from astrocytes. Using the procedure introduced for brain EVs, counting of Vybrant-CM-diI/tGFP particles showed that 89.3% ± 2.2 % are co-labeled particles (Fig. 2C, right green bar), indicating that orbital shaking produced a homogenous population of CD9-tGFP EVs from astrocytes.

EVs from shaken and non-shaken cultures showed identical size distributions (Fig. 2D), but particle numbers increased by up to 20-fold (Fig. 2E), consistent with the co-labeling experiment. Immunoblot analysis showed that astrocyte-derived EVs contained the CD9-tGFP fusion protein, detected exclusively in cells expressing Aldh1l1-driven Cre, confirming astrocyte-specific secretion of CD9-tGFP-labeled EVs (Fig. 2F). Expression of CD9-tGFP did not affect endogenous CD9 levels or canonical EV markers (Flotillin-2, Alix1), indicating that basal EV secretion remained unchanged. In addition to canonical markers, EVs contained Annexin A1 and Arl13b, markers of microvesicles and cilia, respectively [36,37].

### CD9-tGFP Localizes to Astrocytic Processes In Vitro

The detection of Annexin A1 and Arl13b in astrocyte-derived EVs (Fig. 2F) suggested vesicle release from specialized plasma-membrane domains, such as filopodia and cilia. In primary glial cultures from Aldh1l1-Cre; CD9-tGFP^fl/fl^ mice, CD9-tGFP localized to perinuclear vesicles and to plasma membrane protrusions of astrocytes (Fig. 3A), identified as filopodia (arrow in inset). This pattern was consistent with our previous observation that CD9 is enriched in filopodia of CD9-tdTomato-transfected astrocytes [38,39]. Furthermore, Annexin A1 was detected in EVs isolated from primary astrocyte cultures (Fig. 2F), consistent with our earlier findings identifying filopodia-derived EVs as microvesicles generated by budding from the plasma membrane [27,38]. Orbital shaking markedly reduced CD9-tGFP labeling at the plasma membrane, while GFAP staining confirmed preservation of astrocyte integrity (Fig. 3B). Together with the increased number of EVs released under the same conditions, these complementary results demonstrated that mechanical stimulation promotes EV release from astrocytic membrane protrusions, supported by the decreased overall green fluorescence intensity after orbital shaking (Fig. 3C).

**Figure 3.**
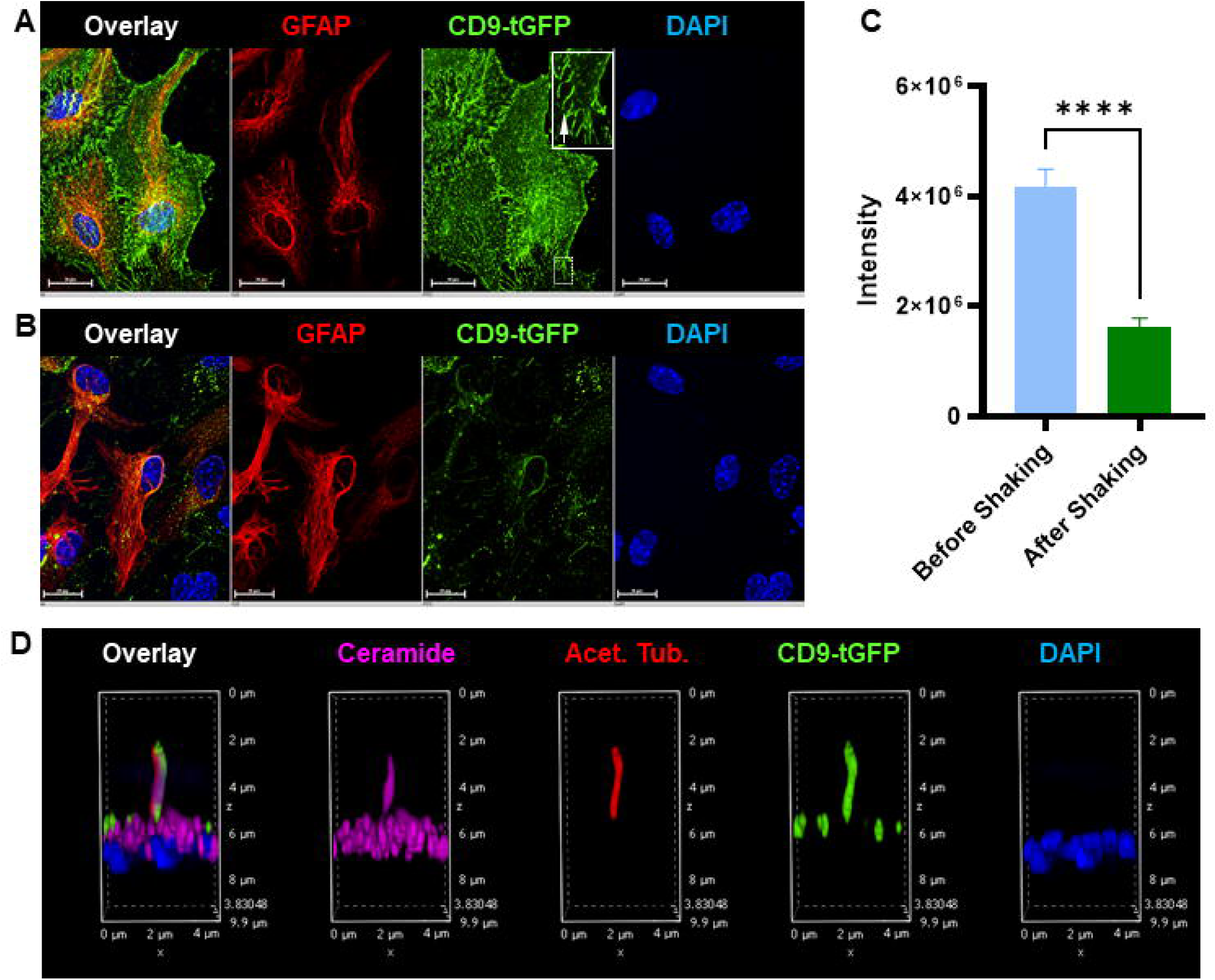
Localization of CD9-tGFP to astrocytic processes *in vitro*. (A) Immunofluorescence imaging of primary-cultured astrocytes from Aldh1l1-Cre; CD9-tGFP^fl/fl^ mice shows CD9-tGFP in perinuclear vesicles and along filopodia-like processes (green) of GFAP-positive astrocytes (red). The inset highlights CD9-tGFP-positive filopodia (arrow). (B) Immunofluorescence imaging of primary astrocytes after orbital shaking. The CD9-tGFP (green) signal is decreased at the plasma membrane, while overall integrity of astrocytes is maintained (GFAP, red). (C) Quantification of the green fluorescence intensity shows that the total green fluorescence largely decreased after orbital shaking (N=10, data are presented as mean ± SEM, t test ****p<0.0001). (D) Immunofluorescence imaging shows CD9-tGFP (green) colocalized with primary cilia labeled by acetylated tubulin (red), and ceramide-enriched compartments (purple), indicating the involvement of CD9-tGFP labeled EVs at astrocytic cilia and ceramide-positive protrusions. Nuclei are counterstained with DAPI (blue).

Likewise, in the astrocytes without shaking, the localization of CD9-tGFP to primary cilia labeled by acetylated tubulin, and co-labeling for ceramide, a sphingolipid critical for microvesicle secretion (Fig. 3D), further supports the hypothesis that CD9-tGFP localizes to plasma membrane protrusions prior to EV secretion from astrocytes.

### CD9-tGFP EVs Secreted by Astrocytes are Transported to Neurons and Capillaries

To determine whether CD9-tGFP localizes to astrocytic processes *in vivo* and to identify the cell types that interact with these processes, we performed immunocytochemistry and fluorescence microscopy using markers for astrocytes (GLAST1, GFAP), microglia (Iba1), capillaries (laminin), and neurons (NeuN, NeuroTrace, Parvalbumin, Calbindin). In the cortex, CD9-tGFP labeling was detected at GFAP-positive peripheral astrocytic processes and adjacent particulate structures (Fig. 4A). consistent with astrocytic membrane protrusions known to closely contact capillaries and neurons. The labeling of peripheral processes frequently extended into lamellipodia-like membrane sheets that co-labeled for GLAST1 (Fig. 4B). While CD9-tGFP expression followed a genotype-dependent increase in homozygous versus heterozygous mice, the spatial distribution pattern remained consistent across genotypes, indicating that expression level did not affect localization of CD9-tGFP (Suppl. Fig. 3A-C). Neither astrocytes nor microglia exhibited strong labeling of their cell bodies (Suppl. Fig. 3D), indicating that CD9-tGFP was primarily localized to astrocytic processes that interface with vascular and neuronal compartments, suggesting these sites as potential targets for astrocyte-derived vesicle signaling. Supplemental Fig. 4A shows stronger CD9-tGFP labeling in highly vascularized regions identified by laminin, a marker for capillaries. At astrocytic endfeet contacting blood vessels, particulate CD9-tGFP was detected on the luminal side of laminin-labeled capillaries, consistent with astrocyte-derived EVs crossing the blood-brain-barrier (Fig. 4C, D and Suppl. Fig. 4B for individual channels).

**Figure 4.**
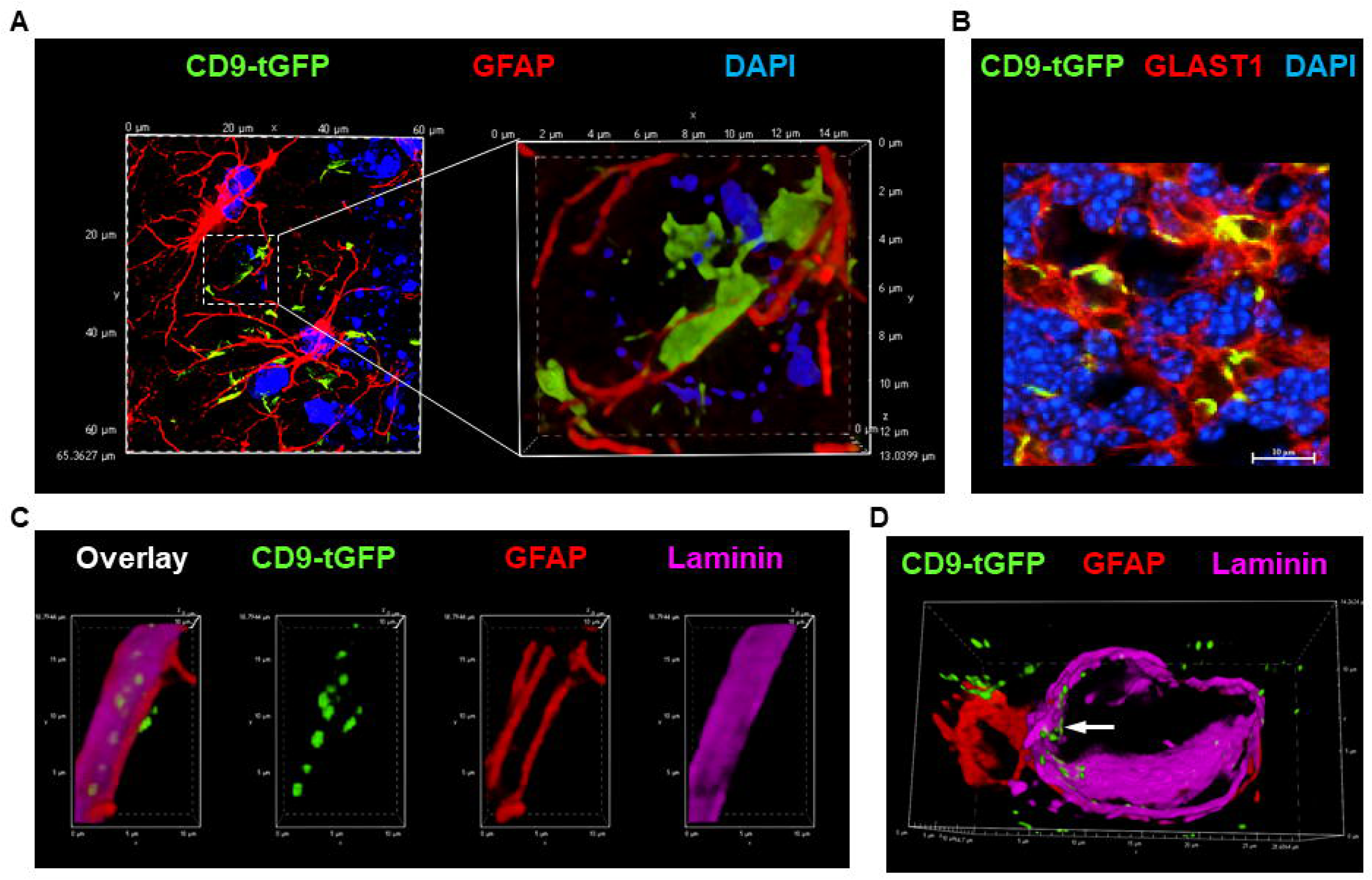
Transport of CD9-tGFP-labeled EVs from astrocytic processes to capillaries *in vivo*. (A, B) CD9-tGFP (green) localizes to lamellipodia-like peripheral astrocytic processes *in vivo*, extending from GFAP-labeled processes (A, red) and co-labeled with GLAST1 (B, red). (C, D) Confocal imaging of brain sections co-labeled for GFAP (red) and laminin (purple) shows CD9-tGFP (green) present on the luminal side of the laminin-positive capillaries in close contact with astrocytic endfeet (arrow). Nuclei are counterstained with DAPI (blue).

To visualize contact sites between CD9-tGFP-labeled astrocytic processes and Purkinje neurons, we performed co-labeling with NeuN and NeuroTrace. The overview (Suppl. Fig. 5A) showed intense NeuN labeling in the pyramidal and granule cell layers of the hippocampus and cerebellum of our mouse model. These NeuN-positive neuronal layers were adjacent to, but did not overlap with, tGFP fluorescence. In the cerebellum, strongly CD9-tGFP-labeled Bergmann glial processes surrounded NeuroTrace-positive neurons morphologically consistent with Purkinje cells (Suppl. Fig. 5B). Co-labeling for calbindin-D28k confirmed the identity of these CD9-tGFP-ensheathed neurons as Purkinje cells (Fig. 5A, and Suppl. Fig. 5C for individual channels). High-resolution images revealed particulate CD9-tGFP signal within calbindin-positive Purkinje somata (Fig. 5B, arrow), indicating these neurons internalized EVs from adjacent astrocytic processes (Suppl. Fig. 6A for individual channels). Parvalbumin immunolabeling of parallel sections delineated the same Purkinje population, highlighting the soma and dendritic arbor ensheathed by CD9-tGFP-positive Bergmann glial processes (Fig. 5C, D, Suppl. Fig. 5D, 6B, 6C for individual channels).

**Figure 5.**
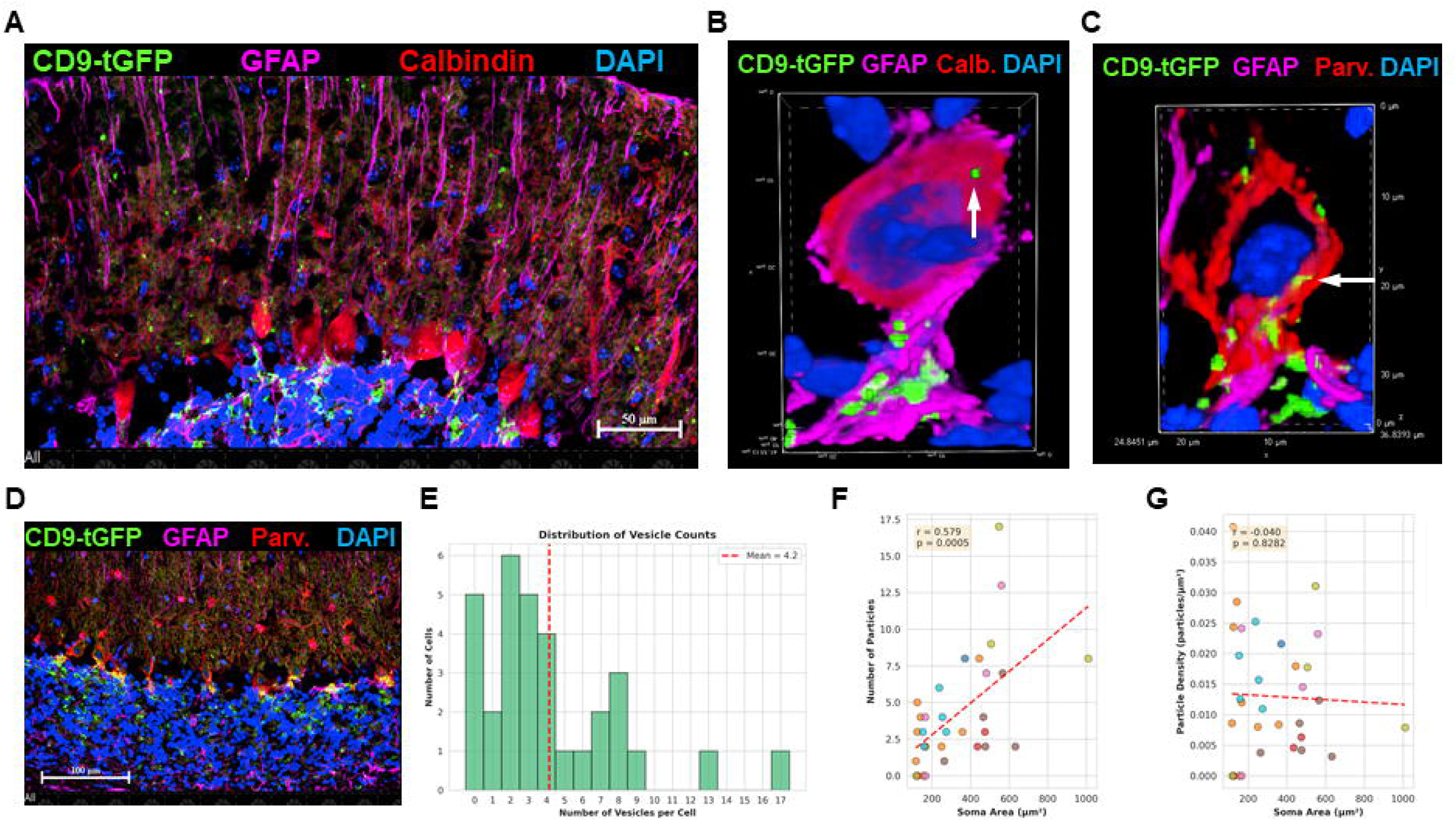
Localization of astrocyte-derived EV cargo in Purkinje neurons. (A) Immunofluorescence imaging of cerebellar sections from Aldh1l1-Cre; CD9-tGFP^fl/fl^ mice labeled for calbindin (red) to identify Purkinje cells colocalized with CD9-tGFP (green), in contact with astrocytic processes (purple). (B) High-resolution imaging shows particulate CD9-tGFP (green) within calbindin-positive Purkinje cells (red). (C, D) Parallel sections immunolabeled for parvalbumin (red) and GFAP (purple) with CD9-tGFP (green) delineate Purkinje neurons and their dendritic arbors ensheathed by CD9-tGFP-positive Bergmann glial processes. (E-G) Quantitation of punctate CD9-tGFP signals inside of parvalbumin-labeled Purkinje neurons using AI-supported image analysis. (E) Distribution of vesicle counts per cell shows a mean of 4.2 CD9-tGFP-positive puncta per Purkinje neuron (n = 3 mice, 7 images, 32 cells, dashed line indicates mean). (F) Particle number positively correlates with Purkinje cell soma size (r = 0.579, p < 0.0005, linear regression shown in red). (G) Particle density (puncta per unit area) shows no significant correlation with soma size (r = -0.040, p = 0.8282 for regression), indicating relatively uniform distribution independent of cell size.

**Figure 6.**
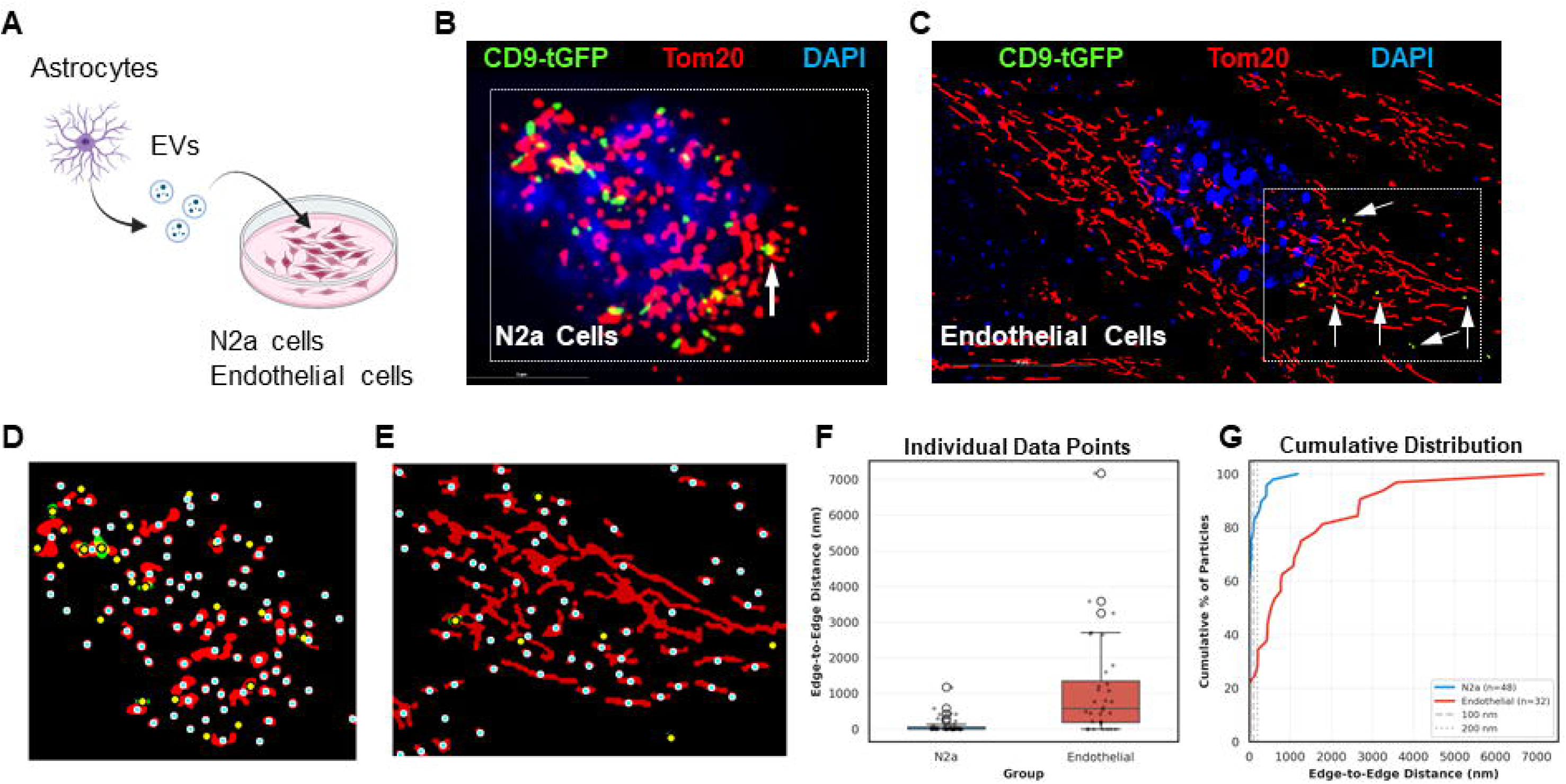
Intracellular distribution of astrocyte-derived EV cargo in N2a cells and primary cultured mouse endothelial cells. (A) Experimental schematic for studying astrocyte-derived EV uptake. Astrocyte-derived CD9-tGFP-positive EVs are isolated from primary cultures and added to N2a cells. (B) Fixed N2a cells 2 h after incubation with CD9-tGFP EVs, immunolabeled for the mitochondrial outer membrane protein Tom20 (red), show CD9-tGFP (green) puncta colocalizing with mitochondria (arrows). C). Fixed primary cultured mouse endothelial cells 2 h after incubation with CD9-tGFP EVs. Labeling as in B. (D-E) AI-supported spatial analysis determining edge-to-edge distance between CD9-tGFP particles and mitochondria in N2a cells (D) and endothelial cells (E). Mitochondria shown as red circles; CD9-tGFP particles color-coded by distance: yellow (≤200 nm), cyan (>500 nm). (F) Quantification shows mean edge-to-edge distance between internalized CD9-tGFP signals and nearest mitochondria; significantly smaller for N2a cells (93 nm) than endothelial cells (1128 nm; Mann-Whitney U, p < 0.0001). Box plots show individual data points overlaid (n=5 independent cell cultures, with 48 N2a particles, 32 endothelial particles, from 5 images each). (G) Cumulative distribution analysis reveals 77% of N2a EVs within 100 nm of mitochondria compared to 22% in endothelial cells, with 60% of N2a EVs in direct contact (0 nm) versus 22% in endothelial cells (Fisher’s exact, p < 0.001).

Using AI, we developed a customized program quantifying the number of EVs taken up by Parvalbumin-labeled Purkinje cells. Critical to this program was application of a binary mask defined by Parvalbumin immunolabeling to exclude CD9-tGFP signals outside of soma boundaries, combined with size filtering to select punctate particles and avoid confusion with extended signal clusters from astrocytic processes. Particles were detected if their centroids fell within the parvalbumin-defined soma mask. Statistical analysis revealed that 67% ± 14% of Purkinje neurons showed internalized EV particles, with a mean of 4.2% ± 0.7% EVs/cell (Fig. 5E). The number of internalized EVs correlated positively with soma size (r = 0.579, p = 0.0005), with larger cells containing more EVs (Fig. 5F). Particle density normalized to soma area showed no correlation with cell size (r = -0.040, p = 0.83) (Fig. 5G), confirming that EV uptake scales proportionally with cell size.

### Astrocyte-Derived EVs are Trafficked to Neuronal Mitochondria In Vitro and In Vivo

The observation that astrocyte-derived CD9-tGFP particles were visible in Purkinje neurons *in vivo* prompted us to monitor EV uptake into live neuronal (N2a) cells. Based on our previous studies showing that EVs can be transported to mitochondria [15,16,27], we incubated N2a cells with astrocyte-derived EVs as shown in Fig. 6A and after fixation, co-labeled cells with Tom20. Figure 6B shows that after 2 h incubation, the CD9-tGFP signal was already detected at mitochondria (arrow). To test if rapid transport of EVs to mitochondria was specific for neuronal cells, we incubated primary cultured brain endothelial cells with astrocyte-derived EVs. Figure 6C shows that while EVs were taken up by endothelial cells, the CD9-tGFP signals were predominantly distributed away from the mitochondrial proximal regions (arrows).

To quantify EV uptake and transport, we used AI to develop a customized program determining the degree of vicinity or apposition of EVs to mitochondria (or any other compartment). Critical to this program was to first determine centroids of objects and then measure the edge-to-edge distance between them (Fig. 6D and E). In addition, the program performed a Monte Carlo simulation to determine if the apposition was distinct from random distribution of objects within a region of interest. Analysis of N2a and endothelial particles revealed that the mean edge-to-edge distance between internalized CD9-tGFP signals and nearest mitochondria was significantly smaller for N2a cells than for endothelial cells (93 nm vs. 1128 nm, Mann-Whitney U, p<0.0001; Fig. 6F and G). Strikingly, 60% of N2a EVs were in direct contact (0 nm) with mitochondria compared to only 22% in endothelial cells (Fisher’s exact p<0.001). Pooled Monte Carlo randomization tests (10,000 iterations) confirmed highly significant enrichment of N2a EVs at mitochondrial surfaces, with 77% within 100 nm versus 8.5% expected by random distribution (p<0.0001), while endothelial cells showed only 22% within 100 nm versus 1.4% expected (p<0.0001). The 3.5-fold difference between cell types was highly significant (p<0.0001) (Suppl. Fig. 7A and B).

Binned distance analysis revealed fundamentally different spatial organizations: in N2a cells, CD9-tGFP-labeled signals showed strong enrichment in close proximity bins (contact and 0-100 nm) with progressive depletion at longer distances, whereas endothelial cells exhibited a bimodal distribution (bimodality coefficient = 0.764) with two distinct populations: one in contact/close proximity (71% of particles, peak at 440 nm) and another distant from mitochondria (29% of particles, peak at 2776 nm) (Suppl. Fig. 7C). Gaussian mixture modeling confirmed that a 2-component model best fit the endothelial distribution (ΔBIC = -17.2), whereas N2a cells showed predominantly unimodal distribution heavily skewed toward close proximity, rather than two genuine biological populations. These findings indicate that in neuronal cells, EV membrane components containing CD9-tGFP are preferentially trafficked to and retained at mitochondria, while in endothelial cells, EVs are either transiently in contact or spatially segregated from mitochondria. A 3D visualization from different angles clearly demonstrates the close vicinity of CD9-tGFP signals to mitochondria in N2a cells (Fig. 7A).

**Figure 7.**
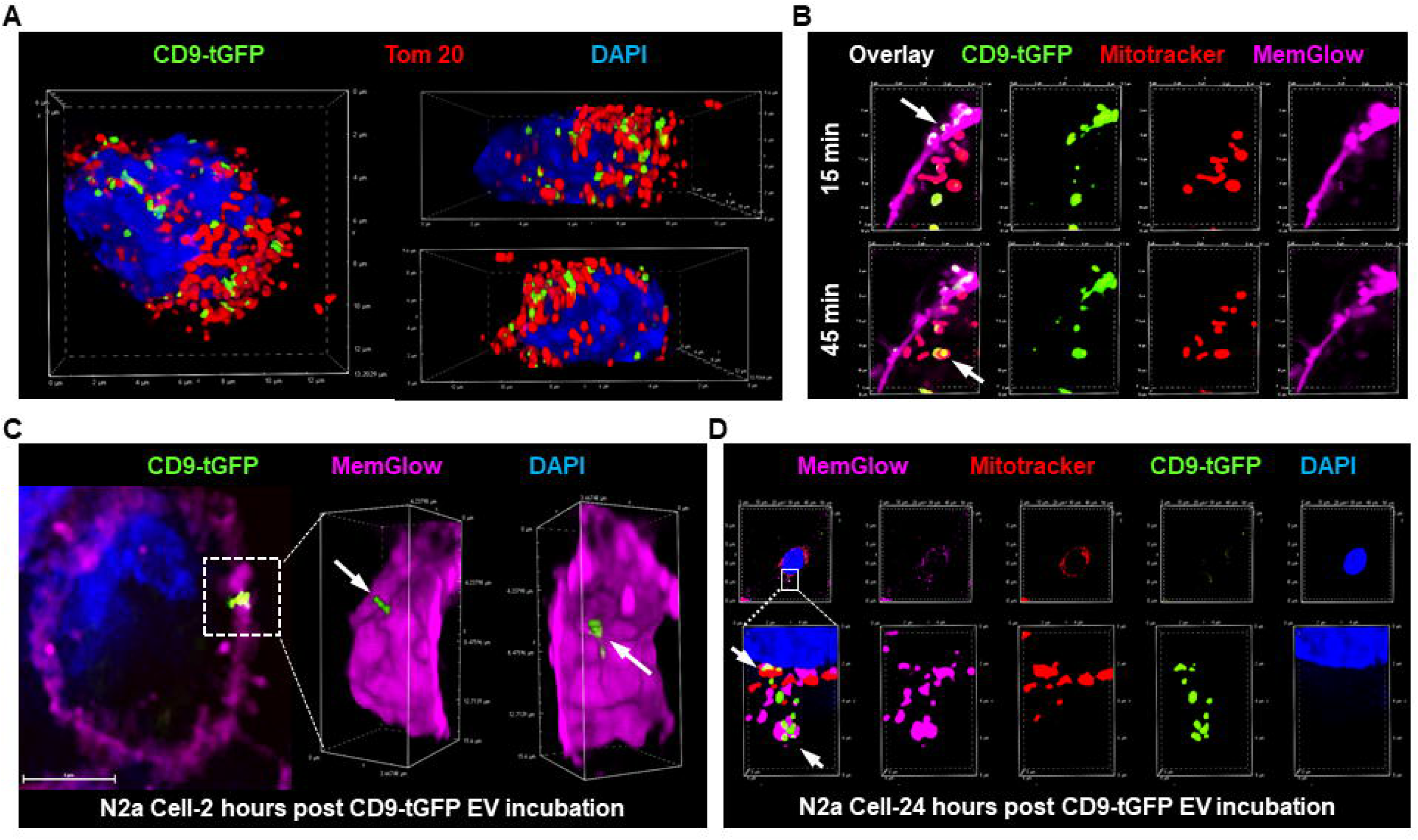
Time-lapse uptake and trafficking of EV cargo in N2a cells. (A) 3D reconstruction of the CD9-tGFP signal from Fig. 6B indicating mitochondria-targeted trafficking of EV cargo in N2a cells. (B) Time-lapse spinning-disk confocal imaging of live N2a cells labeled with MemGlow for plasma membrane (purple) and MitoTracker for mitochondria (red) treated with CD9-tGFP-positive EVs (green). Within 15 min, CD9-tGFP-positive EVs are observed fusing with the plasma membrane (upper panel, arrow), and after an additional 30 min, the CD9-tGFP signal appears as intracellular vesicles and puncta colocalizing with mitochondria in N2a cells (lower panel, arrow). (C) 3D reconstruction of fixed N2a cells showing plasma membrane (purple) fusion after 2 h of incubation with EVs. (D) Different stages of EV cargo internalization and EV cargo transport to mitochondria after 24 h of EV incubation.

To understand uptake kinetics, N2a cells were labeled with fluorescent dyes for the plasma membrane (MemGlow) and mitochondria (MitoTracker), incubated for varying time periods with CD9-tGFP-labeled EVs and cells monitored using spinning disk confocal laser live cell microscopy. As shown in Fig. 7B-upper panel, within 15 minutes after addition to the culture medium, CD9-tGFP-labeled EVs were observed fusing with the plasma membrane (arrow). After an additional 30 minutes (Fig. 7B-lower panel), green fluorescent puncta reappeared as intracellular vesicles or colocalized with mitochondria (arrow), suggesting that CD9-tGFP was transferred either from internalized EVs or directly from the plasma membrane to mitochondria. Plasma membrane fusion, internalization of CD9-tGFP, and mitochondrial targeting were confirmed even 24 h after incubation with EVs in N2a cells (Fig. 7C,D).

To further determine whether trafficking of CD9-tGFP to neuronal mitochondria involves transfer of EV membrane cargo to mitochondria *in vivo*, we isolated synaptic and non-synaptic mitochondria from the brains of Aldh1l1-Cre; CD9-tGFP^fl/fl^ mice using the method shown in Suppl. Fig. 2B. CD9-tGFP fluorescence was detected at the surface of individual mitochondria labeled with MitoTracker (Fig. 8A). Quantitation of mitotracker and tGFP labeling showed that despite the non-synaptic mitochondria fraction being 4-times larger than the synaptic fraction, labeling of synaptic mitochondria was 4-fold higher, indicating that EV cargo was preferentially colocalized with synaptic mitochondria (Fig. 8B). This conclusion was confirmed by immunoblot analysis, showing that consistent with a larger number of mitochondria Tom-20 labeling was higher in the non-synaptic fraction, while aligned with a specific distribution of EV cargo, CD9-tGFP labeling was stronger in the synaptic fraction of mitochondria (Fig. 8C). In contrast, endogenous CD9 was detected in whole-brain lysates but only in trace amounts in the mitochondrial fractions. The detection of CD9-tGFP in purified mitochondria thus supports the conclusion that astrocyte-derived EVs deliver their cargo to neuronal mitochondria. We also verified the *in vivo* transfer of CD9-tGFP EVs to mitochondria in Parvalbumin-positive Purkinje cells using confocal microscopy (Fig. 8D and E, Suppl. Fig. 8 for individual channels) and STED super-resolution microscopy (Fig. 8F) further supporting the *in vivo* targeting of neuronal mitochondria by astrocyte-derived EVs.

**Figure 8.**
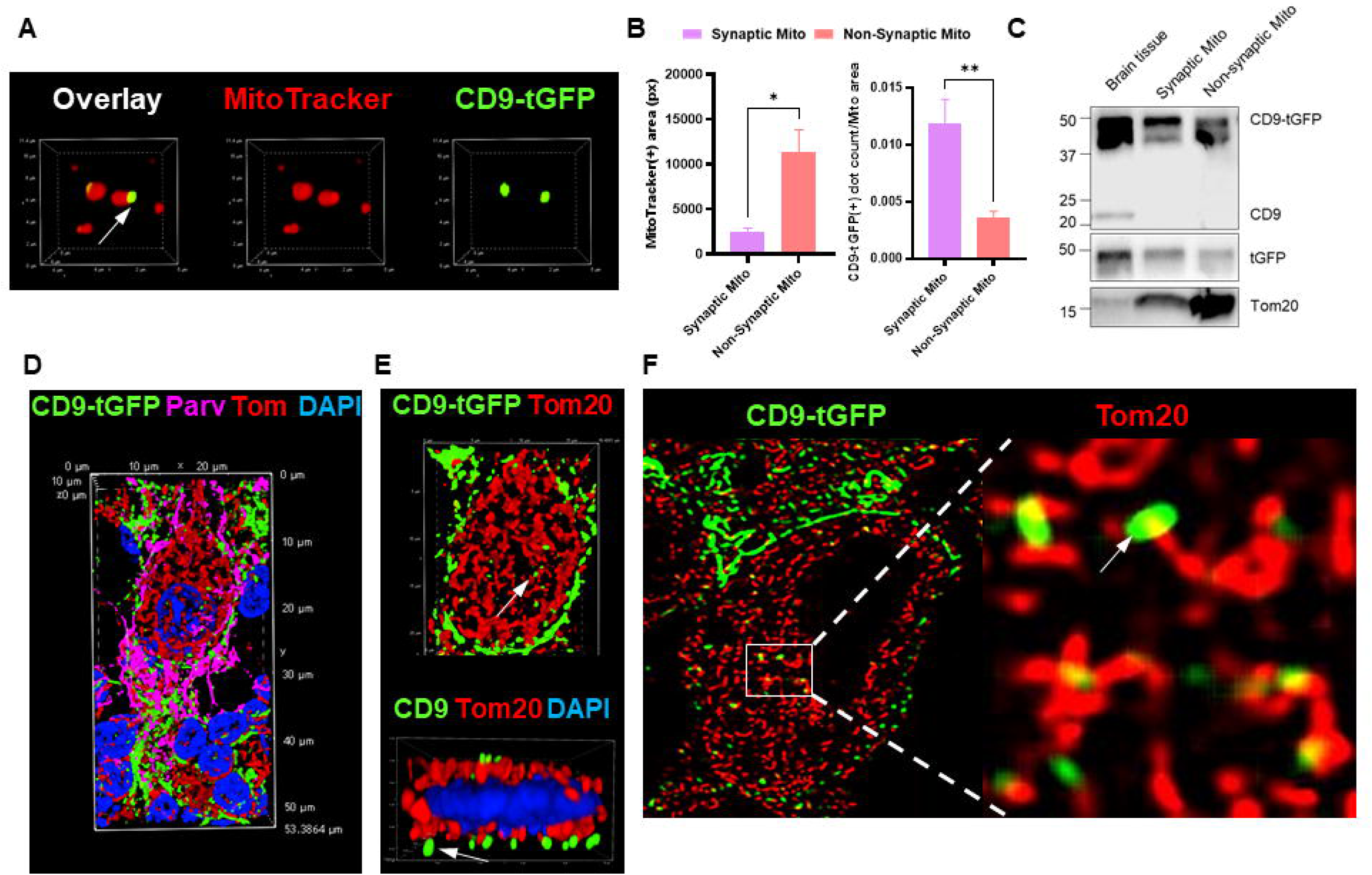
Astrocyte-derived EVs associate with neuronal mitochondria *in vivo*. (A) Immunofluorescence imaging of purified mitochondria from Aldh1l1-Cre; CD9-tGFP^fl/fl^ brains labeled with MitoTracker (red) and colocalized with CD9-tGFP (green), showing CD9-tGFP fluorescence on individual mitochondria. (B) Quantification of the MitoTracker-positive area for mitochondria (left bar graph), and the quantification of the ratio of CD9-tGFP-positive dots to the corresponding mitochondria area (N=5, data are presented as mean ± SEM, t test *p<0.05, **p<0.0001). (C) Immunoblots of brain lysates, synaptic mitochondria, and non-synaptic mitochondria probed for CD9, tGFP, and the mitochondrial marker Tom20. Brain lysates present both endogenous CD9 and EV markers, while mitochondrial fractions present Tom20 only, indicating successful mitochondria isolation. (D, E) Confocal imaging of brain sections shows colocalization of CD9-tGFP fluorescence (green) with Tom20 in neuronal mitochondria (red) of parvalbumin-positive Purkinje neurons (purple). Nuclei are counterstained with DAPI (blue). D shows 3D reconstruction and view from the side. (F) STED super-resolution microscopy imaging of parvalbumin-positive Purkinje neurons confirms the colocalization of CD9-tGFP (green, arrow) with the mitochondrial Tom20 signal (red).

## Discussion

Progress in research on EVs in the CNS has been limited due to the lack of animal models that allow tracking of EVs from their cellular origin to their targets *in vivo* [12,40]. Lipophilic dyes, often used to label EVs are prone to non-specific transfer, dye aggregation and diffusion causing false-positive signals and limited subcellular resolution [41]. CD63 and CD81 reporters primarily label EV subsets enriched in these two tetraspanins. Neckles et al. developed the TIGER (CD9-tGFP) reporter mouse as a Cre-dependent EV-labeling tool, but astrocyte-specific applications have not yet been reported. The CD9-tGFP model expands EV coverage by labeling vesicles carrying CD9, thereby capturing a broader spectrum that includes EV populations predominantly released via plasma membrane budding. In this paper, we present the first astrocyte-specific implementation by crossing TIGER with Aldh1l1-Cre to generate Aldh1l1-Cre; CD9-tGFP^fl/fl^ mice. This model enables highly selective labeling and visualization of astrocyte-derived CD9-tGFP-positive EVs both *in vitro* and *in vivo*. Beyond showing trafficking of EV-derived cargo to neuronal mitochondria using high-resolution fluorescence microscopy, we isolated synaptic and non-synaptic mitochondria and detected CD9-tGFP in both fractions. These results demonstrate that astrocyte-derived CD9-positive EVs are taken up by neurons and their cargo trafficked to mitochondria, a process not previously been documented *in vivo*.

Previously, our lab proposed that EVs can be viewed as “mobile rafts” that export specific lipid signaling platforms into the extracellular space [42]. EV biogenesis is tightly linked to sphingolipid metabolism and transport, as our lab previously showed that EV formation and secretion is regulated by ceramide transfer protein (CERT) and sphingomyelinases (SMases)-mediated ceramide release, both establishing the important involvement of ceramide in EV sphingolipid composition, formation, and function [38]. The association of ceramide with tetraspanins such as CD9 in ceramide-rich platforms (CRPs) is of particular importance since both induce membrane curvature, a prerequisite to vesicle formation. We reported for the first time that ceramide is specifically enriched in CD9-positive EVs released from astrocytic filopodia, an observation that may relate to the increased EV production by astrocytes under pathological stress conditions such as Alzheimer’s disease [27].

These findings show that the Aldh1l1-Cre; CD9-tGFP^fl/fl^ model is well suited for visualizing astrocyte-derived CD9-positive EV secretion, including stress-responsive release and lipid-driven (ceramide-mediated) release. Previous mouse models for *in vivo* EV tracking were based on CD63 or CD81 [43,44], which label specific EV subsets but may not adequately represent the ceramide-rich, stress-responsive CD9-positive EV population we had characterized. Previous studies showed that CD9 preferentially localized to plasma membrane budding sites and microvesicles [25–27]. In the current study, we show that CD9-tGFP is enriched in filopodia and cilia of primary cultured astrocytes, consistent with our finding that CD9-tGFP-positive EVs contain Annexin A1 and Arl13b, markers of microvesicles and cilia, respectively, indicating a CD9-positive EV population budding from astrocytic filopodia and cilia. Additionally, the co-labeling of CD9-tGFP with ceramide at these sites further supports the hypothesis that Arl13b-positive EVs originated from cilia. In brain sections from the reporter mouse model, we found that CD9-tGFP fluorescence colocalizes with astrocytic processes, often in endfeet associated with laminin-labeled capillaries or astrocytic membranes engulfing neurons. Notably, fluorescence was also found on the luminal side of capillaries suggesting that astrocytes secrete EVs into the blood stream, most likely through endothelial cells. We also found particulate CD9-tGFP fluorescence in neurons in the mouse brain, demonstrating that the Aldh1l1-Cre; CD9-tGFP^fl/fl^ mouse is valuable model for *in vivo* tracking of astrocyte-derived EVs. To quantify uptake of astrocyte-derived EVs into neurons *in vivo*, we used AI to develop a novel analysis platform specifically customized to our reporter mouse. The analysis revealed that in the cerebellum, internalized CD9-tGFP signals were correlated with Purkinje cell area, suggesting the uptake mechanism is not saturated and likely dependent on the plasma membrane surface, which is consistent with previously published uptake of EVs or their cargo by endocytosis [45–48].

A major knowledge gap in EV research is whether internalized EVs are primarily routed to degradative pathways or can instead functionally integrate with intracellular organelles [13,45,49]. Our observation that astrocyte-derived CD9-tGFP fluorescence colocalizes with neurons *in vivo* prompted us to use CD9-tGFP-labeled EVs from primary cultured astrocytes for live cell uptake studies with neuronal (N2a) cells *in vitro*. Structurally, CD9 is a type III transmembrane protein with the N- and C-terminal sites pointing toward the cytosol [50]. While the plasma membrane buds outward to form microvesicles, the C-terminal domain of CD9 consequently is within the inner lumen of the vesicles. As the tGFP moiety is tagged to the C-terminus of CD9 tetraspanin [17], the CD9-tGFP domain resides at the luminal side of CD9-positive EVs released from astrocytes. However, if CD9-tGFP EVs or their membrane-derived cargos fuse with the plasma membrane or with any intracellular compartment, the tGFP moiety will again point to the cytosol.

Using spinning disk confocal with super resolution module, our data suggest that uptake of primary astrocyte-derived EVs begins with their fusion to the N2a plasma membrane. However, CD9-tGFP fluorescence remains localized and reappears as punctate labeling in intracellular vesicles and at mitochondria, indicating that part of the EV membrane remains intact and undergoes *in situ* internalization. This EV membrane identity aligns with the role of EVs as “membrane presentation platforms” [51] or “mobile rafts” [39,52,53], and is consistent with CRPs and associated tetraspanins maintaining structural organization after membrane integration. At this point, it is not clear whether rapid transport to mitochondria occurs directly from the plasma membrane or via endocytosis. It should be noted that in contrast to transport of the exogenously derived CD9-tGFP signal in N2a cells, donor astrocytes do not show colocalization of CD9-tGFP with mitochondria (not shown), consistent with the distribution of endogenous CD9 to the endosomal compartment and the plasma membrane. The association of EVs with neuronal mitochondria was confirmed by immunofluorescence microscopy of mitochondria isolated from brain. The appearance of fluorescent buds on the surface of MitoTracker-labeled mitochondria is consistent with the topology of CD9-tGFP, which positions the C-terminal tGFP moiety toward the cytosol. The association or fusion of CD9-tGFP-positive EVs with mitochondria is further supported by immunoblots confirming the expression of CD9-tGFP in synaptic and non-synaptic mitochondrial fractions from the brain of our mouse model. Confocal and high-resolution STED microscopy imaging are consistent with these findings, suggesting that EVs are functionally delivered by membrane fusion, facilitating the delivery of lipid and protein cargos into recipient organelles, which is a process potentially critical for the mitochondrial regulation observed in aging and neurodegeneration [23,54].

Importantly, the distribution of internalized CD9-tGFP differed between neuronal (N2a) and primary cultured endothelial cells. Using AI, we developed a second analytical platform customized to quantify the apposition, contact, and overlap of CD9-tGFP signals with intracellular compartments and organelles, particularly mitochondria. In addition, we added a novel Monte Carlo-based randomization method to test if the distribution of internalized CD9-tGFP signals was distinct from random and different in N2a and endothelial cells exposed to astrocyte-derived EVs. The analysis showed that in N2a cells transport of EVs or EV-derived cargo is primarily (>80%) directed to mitochondria, while data from endothelial cells are best fitted by proposing a bimodal distribution of EVs with a portion (<25%) being trafficked to mitochondria, at least temporarily, while the remaining portion is transported to a different compartment.

This distinct distribution of signals suggests that specific cell types affect the fate of EVs in different modalities and cell environment. The specificity of the recipient cell types and destination organelles indicates that astrocyte-derived EVs carry information to mediate communication between astrocytes and neurons. Previous studies have demonstrated that astrocyte-derived EVs can transfer neuroprotective cargo to neurons [55,56], including cargo derived from astrocytic mitochondria [57,58]. However, many studies utilized lipophilic dyes, a method often limited to visualizing general cellular uptake of EVs by neurons rather than resolving precise intracellular location and tracking [59]. Additionally, there is evidence supporting the bi-directional communication between neurons and astrocytes through EVs [44,60]. Nevertheless, the specific fates of astrocytic EVs at the organelle level within recipient neurons have not been fully characterized, primarily due to the lack of *in vivo* tools for cell type and tissue-specific tracking of EVs. Our mouse model is effectively providing evidence tracking the organelle- and cell type-specific integration of EVs both *in vitro* and *in vivo*, resolving these ambiguities associated with non-specific dye transfer and defining the precise organelle destination. Currently, our data cannot distinguish between the physical association or fusion of EV-derived cargo with the outer membrane neuronal mitochondria. The initial fusion of CD9-tGFP signals with the plasma membrane followed by apposition or partial overlap of CD9-tGFP signals with mitochondria indicates that EV-derived membrane components are internalized mainly via endocytosis and subsequently transported to mitochondria, likely entertaining endosomal contact sites to facilitate the intercellular transfer of these astrocytic EV cargos to mitochondria [48,61]. Whether the EV cargo is functionally integrated with neuronal mitochondria and why and how the EV transport pathway differs between neuronal and endothelial cells will be investigated in future studies.

The discovery that astrocyte-derived EVs target neuronal mitochondria places our mouse model and findings within the broader landscape of astrocyte-neuron communication in the CNS. Previous studies have shown that the transfer of EVs containing astrocytic mitochondria or mitochondrial compartments to other cell types, including endothelial cells and neurons, increases with aging, while astrocyte-released EVs can provide neuroprotection in neurodegenerative conditions [54,62]. In such conditions, astrocyte-derived mitochondrial support appears to be neuroprotective, enhancing synaptogenesis and metabolic resilience [59,63]. However, these previous studies did not report transport of EVs to mitochondria of the recipient cells. In contrast, our lab previously demonstrated that astrocytic EVs can exert deleterious effects on neurons by transporting toxic cargo such as ceramide and Aβ to mitochondria, leading to mitochondrial dysfunction, neurotoxicity, and neuronal cell death [14,23,27]. Using our mouse model, these questions about EV function can now be addressed *in vivo* to define the physiological and pathophysiological roles of astrocyte-to-neuron EV transfer. This is particularly relevant in disease contexts, including neurodegenerative disorders, epilepsy, and HIV-associated neurocognitive disorders, where astrocyte-derived EVs have already been implicated in both neuroprotective and neurotoxic processes [8,9,11,64–66].

Notably, in the cerebellum, astrocyte-derived EVs were selectively transported to mitochondria of Calbindin- and Parvalbumin-positive neurons, consistent with the well-established functional coupling between Purkinje neurons and Bergmann glia. Although correlative, this pattern is compatible with preferential targeting of neuronal populations with high mitochondrial ATP demand imposed by inhibitory synaptic activity and astrocyte-dependent glutamate handling. Our astrocyte-specific EV reporter mouse provides an *in vivo* model which can be applied to study how EVs target distinct cell types and organelles across different disease stages, brain regions, or in response to therapeutic intervention. Increasing numbers of studies have used EVs as delivery vehicles for membrane-associated therapeutic proteins or engineered cargo [6,67]. The ability to visualize astrocyte-derived EV release and mitochondrial targeting *in vivo* will facilitate preclinical experiments of interventions that modulate EV biogenesis, redirect EV trafficking, or enrich EVs with neuroprotective and mitochondria-associated cargo, thereby helping to resolve the functional consequence of astrocytic EV-neuronal mitochondrial engagement.

In summary, our Aldh1l1-Cre; CD9-tGFP^fl/fl^ mouse model overcomes a major barrier in research on EVs in the CNS by enabling cell-type specific tracking of astrocyte-derived EVs *in vivo*. Our data show that these EVs arise from specialized plasma-membrane domains, reach vascular and neuronal compartments, and deliver membrane cargo to neuronal mitochondria both *in vitro* and *in vivo*. Our data also show that uptake or transport pathways are specific for different cell types. This establishes a previously unaddressed mechanism of astrocyte-neuron communication and provides a robust tool for future studies on how astrocytic EV-mediated mitochondrial targeting contributes to both neuroprotection and neurodegeneration.

## Acknowledgments

This work was funded by grants from the National Institutes of Health R21AG078601 and RF1AG078338, and the Department of Veterans Affairs 1I01BX007282. We thank Dr. David Feliciano, Clemson University, for consultation with the TIGER mouse and Dr. Patrick Sullivan, University of Kentucky, for help with isolation of mitochondria. We thank the Light Microscopy Core at University of Kentucky for their instruments and assistance with confocal and super resolution imaging. We are grateful for institutional support by the Department of Physiology (Chair Dr. Alan Daugherty).

