## Supplemental Data for "A novel reporter mouse for astrocyte-derived extracellular vesicles reveals trafficking of cargo to neuronal mitochondria"

**Supplemental Figures**

**Supplemental Figure 1**

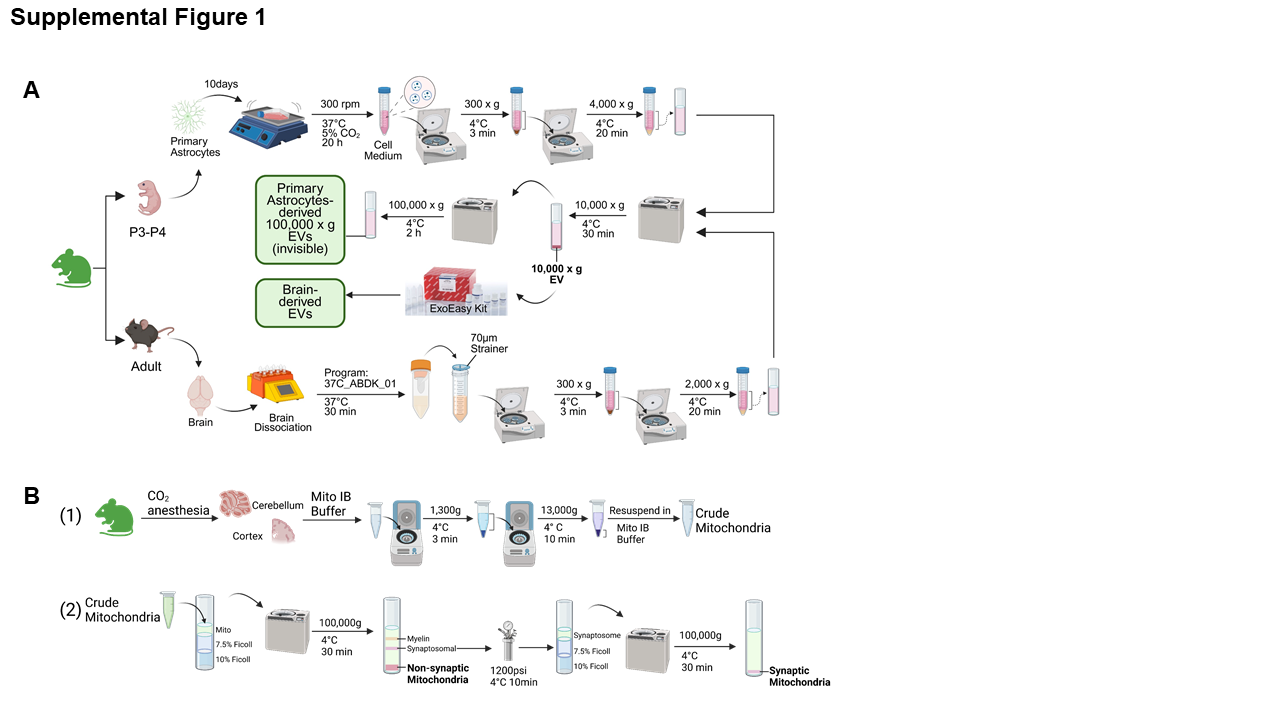

**Workflow for isolation and characterization of astrocyte-derived EVs and mitochondria.**

(A) Schematic of the protocol used to isolate EVs derived from primary astrocyte cultures and brain, respectively, from Aldh1l1-Cre; CD9-tGFP^fl/fl^ mice. (B) Schematic of the protocol used to isolate synaptic and non-synaptic mitochondria from mouse brain.

**Supplemental Figure 2**
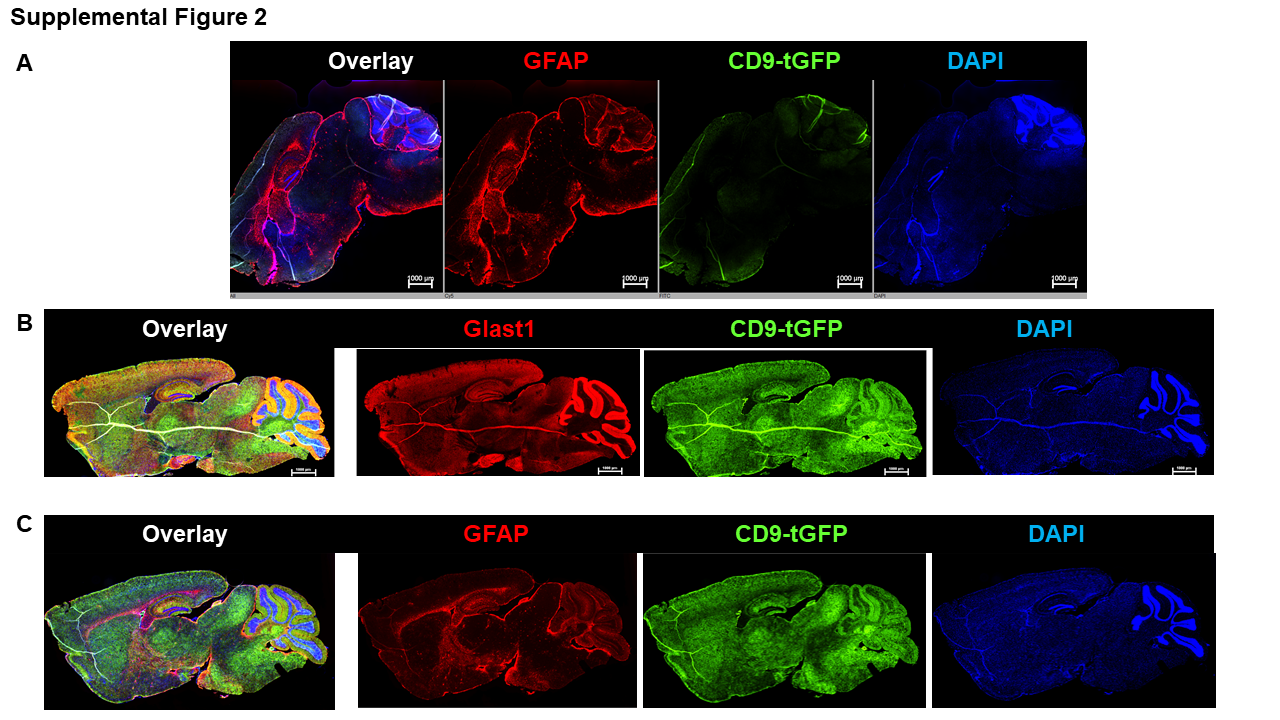

**Regional distribution and specificity of CD9-tGFP expression.**

(A) Whole-brain fluorescence scan of a CD9-tGFP^fl/fl^ mouse lacking Aldh1l1-Cre (no Cre) showing only weak background autofluorescence (green channel). (B) Immunofluorescence imaging of overlay, GLAST1 (red), CD9-tGFP (green), and DAPI (blue) in the brain overview of Aldh1l1-Cre; CD9-tGFP^fl/fl^ mice, corresponding to Figure 1E. (C) Immunofluorescence imaging of overlays, GFAP (red), CD9-tGFP (green), and DAPI (blue) in the parallel section, corresponding to Figure 1F.

**Supplemental Figure 3**
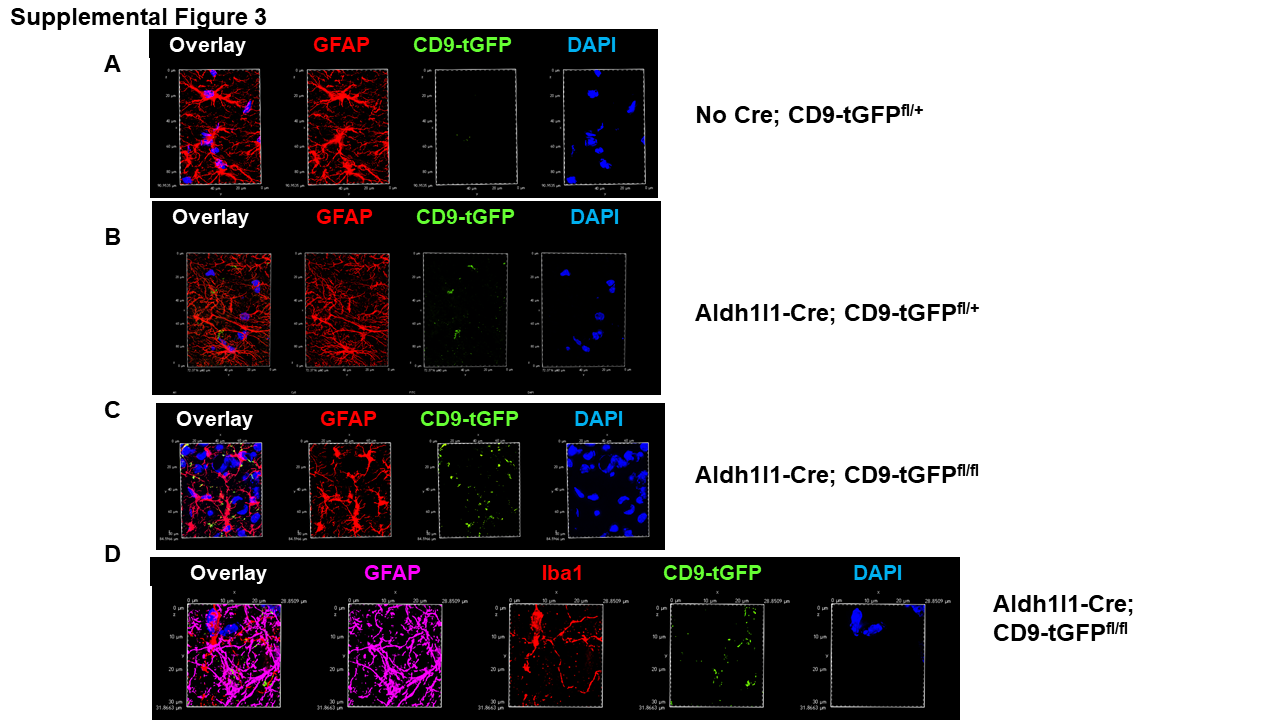

**Expression of CD9-tGFP independence on zygosity and cell type.**

(A, B, C) CD9-tGFP^fl/fl^ mice lacking Aldh1l1-Cre (No Cre) show minimal CD9-tGFP signal (A, green) in GFAP-positive astrocytes (A, red). Heterozygous Aldh1l1-Cre; CD9-tGFP^fl/+^ mice exhibit moderate astrocytic CD9-tGFP fluorescence (B, green) colocalizing with GFAP (B, red). Homozygous Aldh1l1-Cre; CD9-tGFP^fl/fl^ mice show stronger CD9-tGFP expression (C, green) in GFAP-positive astrocytes (C, red) with a similar distribution pattern (A, B, C, red) indicating that increased reporter gene dosage enhances signal intensity without altering localization. (D) Immunofluorescence imaging of brain sections of Aldh1l1-Cre; CD9-tGFP^fl/fl^ mice co-labeled with CD9-tGFP (green), GFAP (purple), and Iba1 (red) demonstrates that neither astrocytes nor microglia exhibit strong CD9-tGFP labeling in cell bodies, supporting a predominant localization of CD9-tGFP to astrocytic processes interfacing with vascular and neuronal compartments. Nuclei are counterstained with DAPI (blue).

**Supplemental Figure 4**

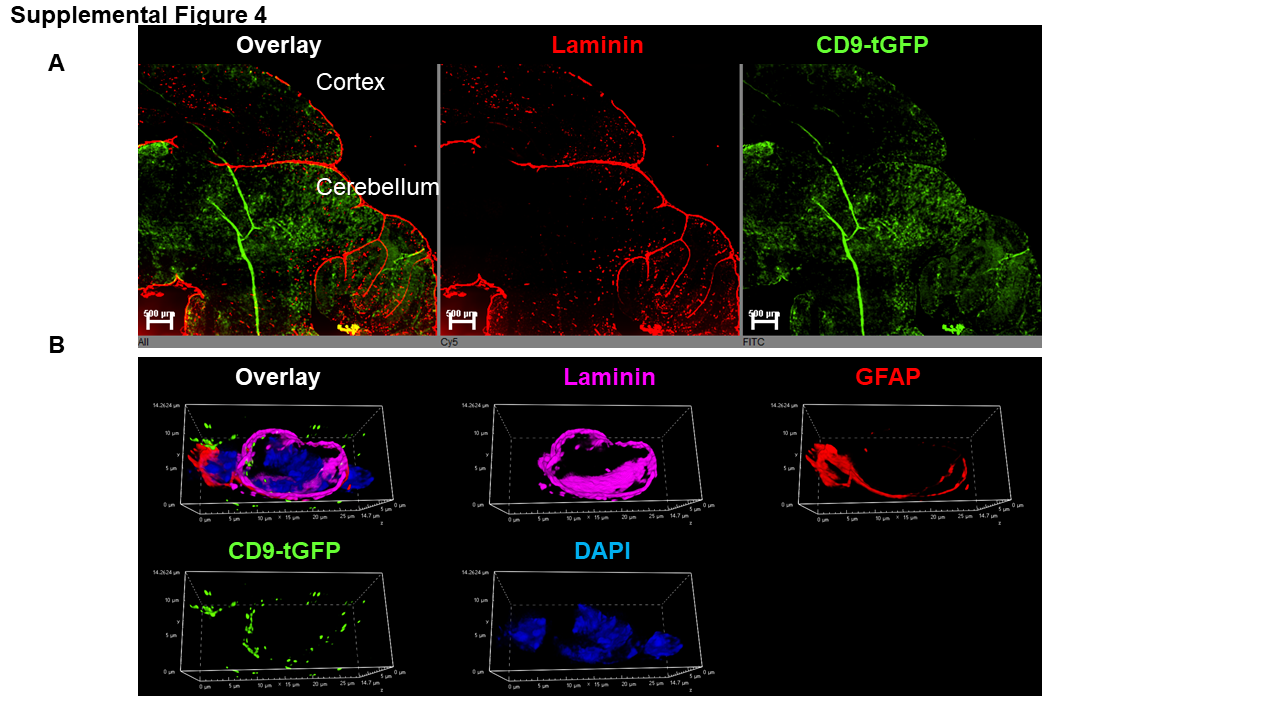

**Expression of CD9-tGFP in highly vascularized brain regions.**

(A) Immunofluorescence imaging of cortex and cerebellum from Aldh1l1-Cre; CD9-tGFP^fl/fl^ mice labeled with laminin (red). CD9-tGFP fluorescence (green) at astrocytic endfeet is prominent in highly vascularized regions. (B) High-resolution imaging overlays of laminin (purple) and CD9-tGFP (green) reveal particulate CD9-tGFP on the luminal side of capillary walls in contact with GFAP-positive astrocytes (red).

**Supplemental Figure 5**

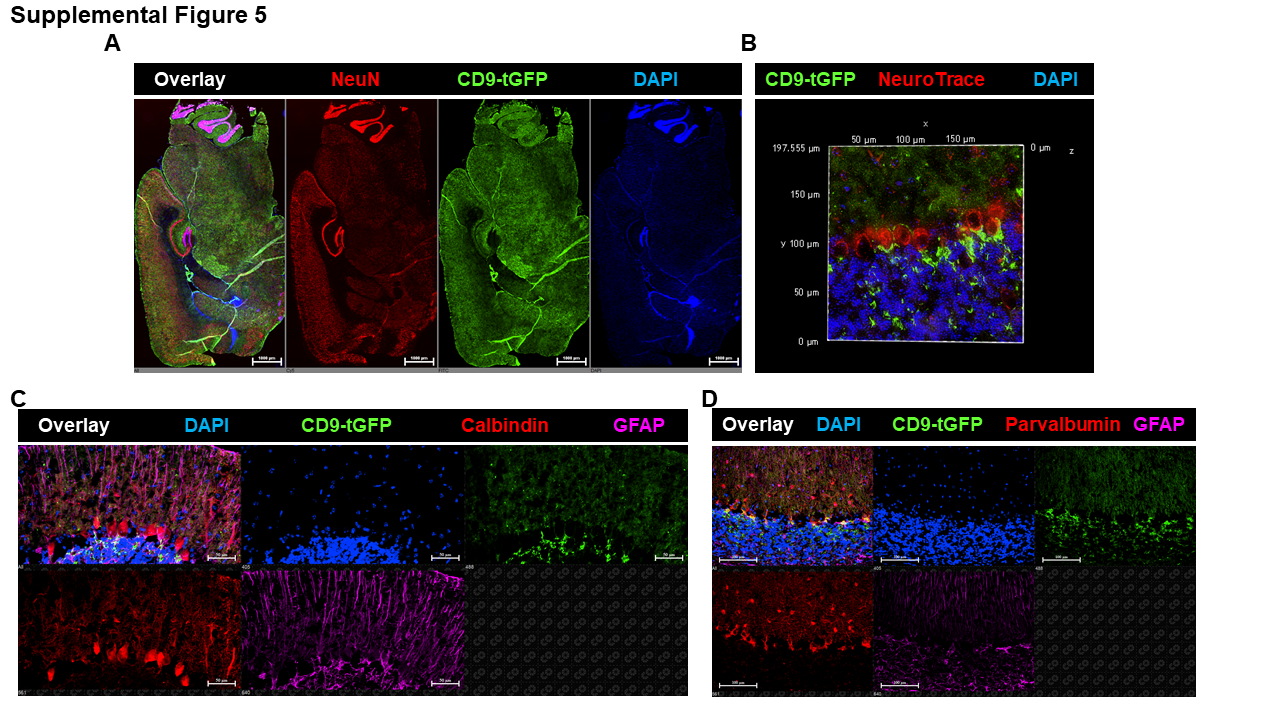

**Interaction of CD9-tGFP-labeled astrocytic processes with neurons.**

(A) Overview of brain imaging from Aldh1l1-Cre; CD9-tGFP^fl/fl^ mice. NeuN-positive neurons (red), particularly at the hippocampus and cerebellum, are adjacent to astrocytic CD9-tGFP fluorescence (green). (B) Immunofluorescence imaging of the cerebellum region labeled with NeuroTrace (red) surrounded by CD9-tGFP-labeled Bergmann glial processes (green). (C) Individual channels show CD9-tGFP fluorescence (green) in calbindin-positive Purkinje somata (red). (D) Parallel sections labeled for parvalbumin-positive Purkinje neurons (red) and associated interneurons ensheathed by CD9-tGFP-positive astrocytic processes (green). Nuclei are counterstained with DAPI (blue).

**Supplemental Figure 6**

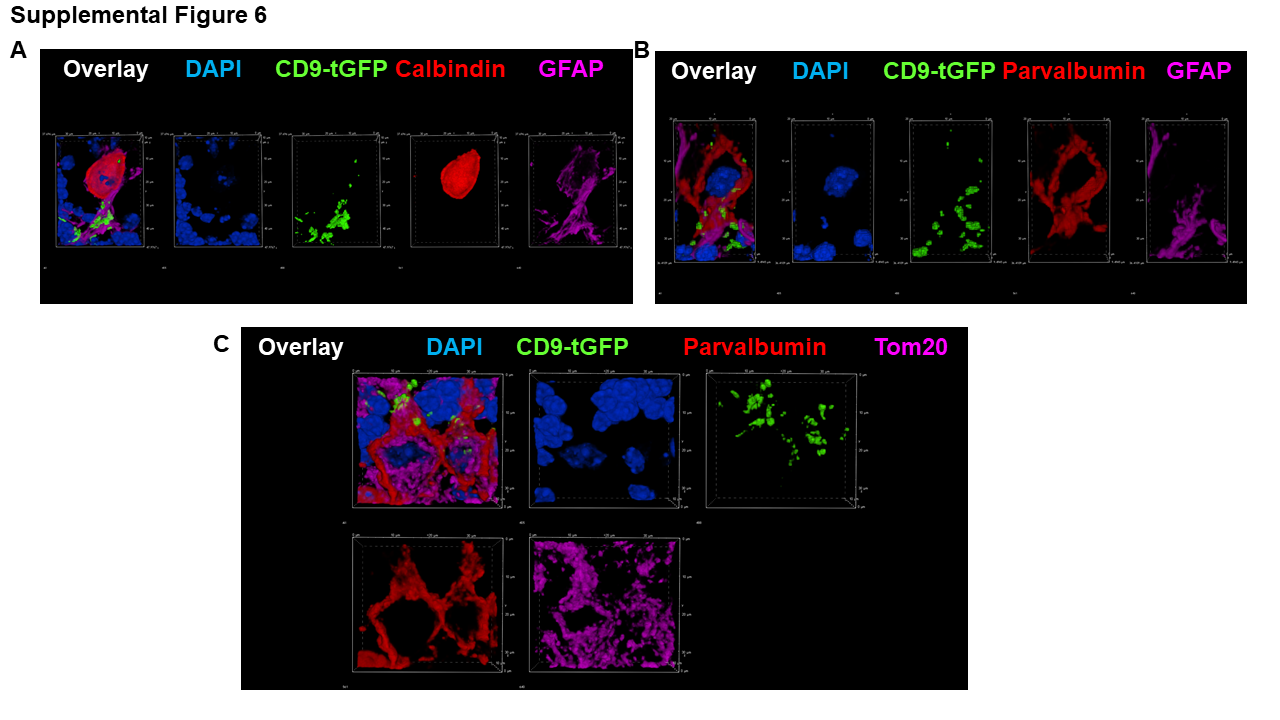

**Uptake and apposition of CD9-tGFP to mitochondria in neurons in vivo.**

(A, B) High-resolution imaging of individual channels showing particulate CD9-tGFP (green) localize in calbindin-positive Purkinje neurons (A, red) and associate with the membrane of parvalbumin-positive Purkinje neurons (B, red). These neurons are in contact with adjacent GFAP-positive astrocytes (purple). (C) High-resolution imaging shows the apposition of CD9-tGFP (green) to mitochondria (Tom20, purple) of parvalbumin-positive Purkinje neurons (red). Nuclei are counterstained with DAPI (blue) throughout.

**Supplemental Figure 7**

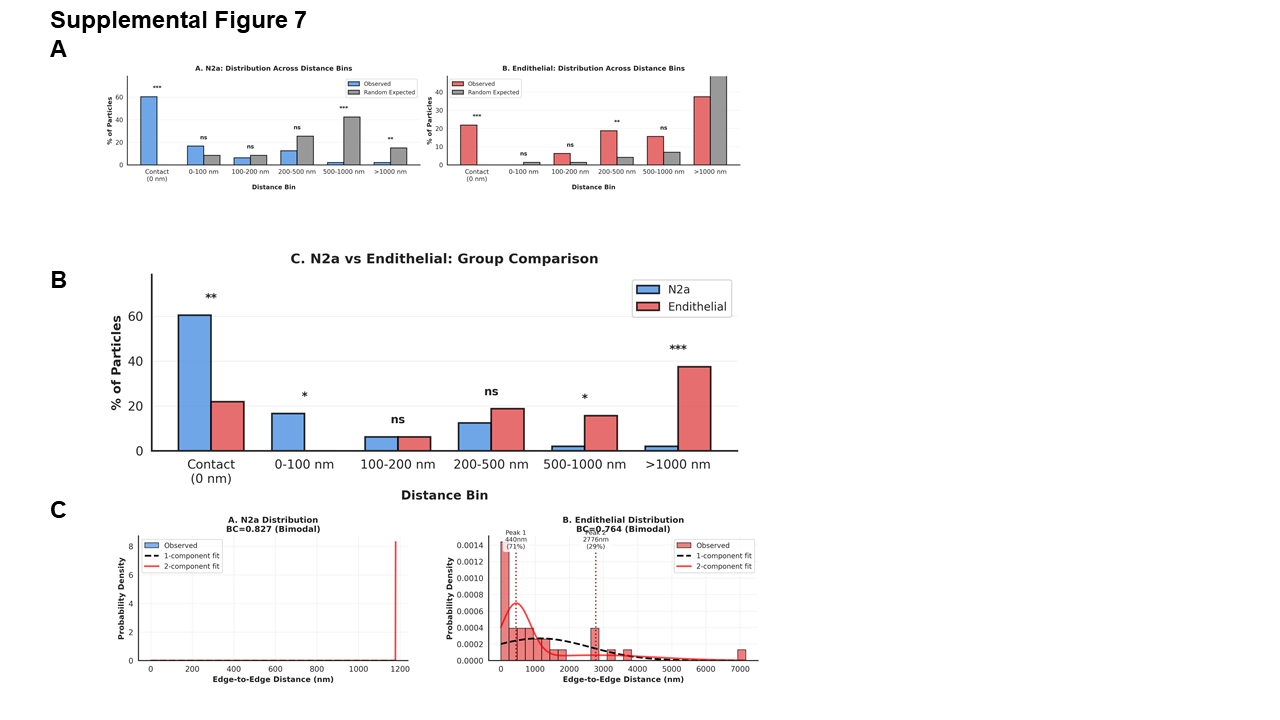

**Apposition analysis of CD9-tGFP signals in N2a cells and primary cultured endothelial cells**

(A) Monte Carlo randomization analysis comparing observed versus expected random

distribution of CD9-tGFP particles across distance bins. Left: N2a cells show highly significant enrichment at contact (0 nm, 60% observed vs. <5% expected, ***) and 0-100 nm bins (17% vs. <5%, ***), with depletion at longer distances. Right: Endothelial cells show significant enrichment at contact (22% observed vs. <5% expected, ***) but more particles at distant bins (>1000 nm: 37% vs. ~40% expected, ***), indicating spatial segregation from mitochondria. Statistical significance determined by 10,000 Monte Carlo iterations comparing observed distribution to random placement within the same region of interest. *p < 0.05, **p < 0.01, ***p < 0.001, ns = not significant. (B) Direct comparison of particle distribution across distance bins between N2a (blue) and endothelial (red) cells. N2a cells show significantly higher percentage at contact (60% vs. 22%, **) and 0-100 nm (17% vs. 3%, *), while endothelial cells show significantly higher percentage at >1000 nm (37% vs. 10%, ***). Chi-square or Fisher's exact test for each bin. *p < 0.05, **p < 0.01, ***p < 0.001, ns = not significant. (C) Bimodality analysis using Gaussian mixture modeling. Left: N2a distribution formally meets the bimodality criterion (BC = 0.827, exceeding the 0.555 threshold) and GMM favors a 2-component fit (ΔBIC = −29.4); however, the 2-component model reflects a dominant near-contact population (98% of particles, peak at ~70 nm) with a single outlier particle (~1179 nm), rather than two genuine biological populations. Histogram (bars) overlaid with 1-component fit (dashed line) and 2-component fit (red line). Therefore, it is treated as predominantly unimodal pattern heavily skewed toward close proximity (contact/0-100 nm). Histogram (bars) overlaid with 1-component fit (dashed line) and 2-component fit (red line). Right: Endothelial distribution exhibits significant bimodality (BC = 0.764, exceeds threshold) with two distinct populations: Peak 1 at 440 nm (71% of particles, close proximity) and Peak 2 at 2776 nm (29% of particles, distant). Gaussian mixture modeling with Bayesian Information Criterion confirms 2-component model best fits endothelial data (ΔBIC = -17.2, favoring bimodal model), while N2a shows predominantly unimodal distribution.

**Supplemental Figure 8**

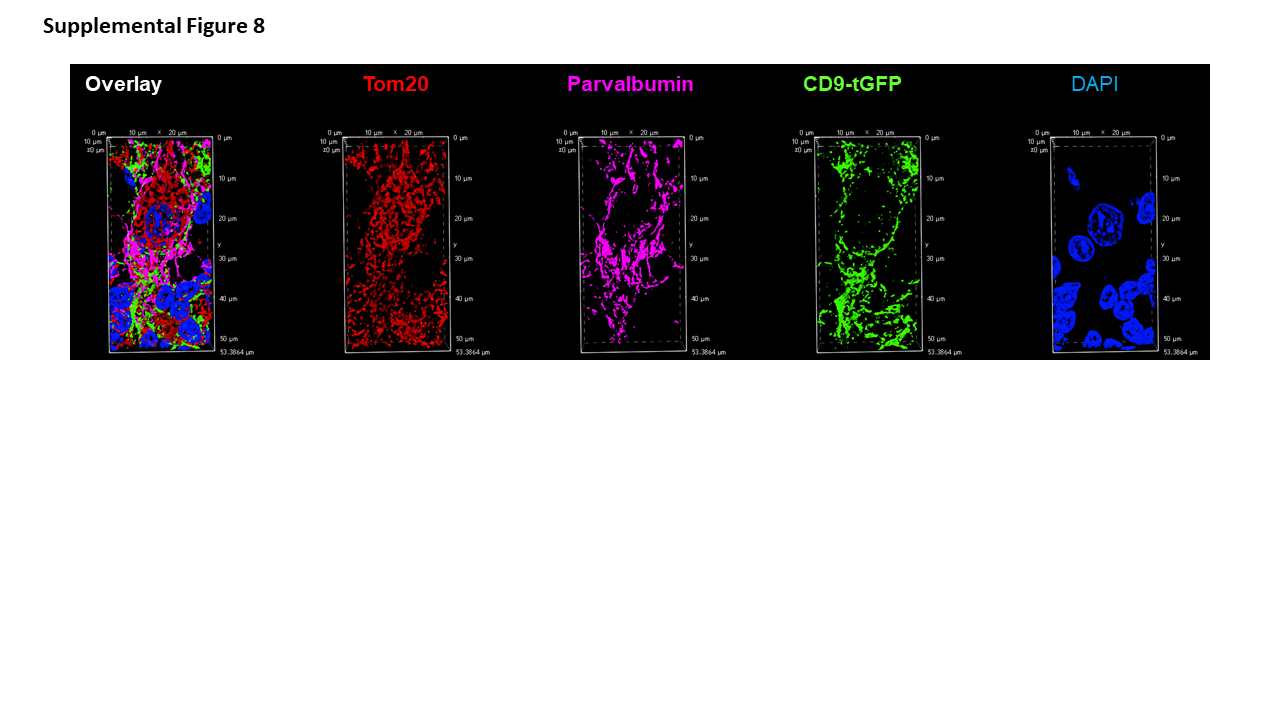

**Individual channels of Fig. 8C**

Confocal imaging of brain sections shows colocalization of CD9-tGFP fluorescence (green) with Tom20 in neuronal mitochondria (red) of parvalbumin-positive Purkinje neurons (purple). Nuclei are counterstained with DAPI (blue).
